# A time and single-cell resolved model of hematopoiesis

**DOI:** 10.1101/2022.09.07.506735

**Authors:** Iwo Kucinski, Joana Campos, Melania Barile, Francesco Severi, Natacha Bohin, Pedro N Moreira, Lewis Allen, Hannah Lawson, Myriam L R Haltalli, Sarah J Kinston, Dónal O’Carroll, Kamil R Kranc, Berthold Göttgens

## Abstract

The paradigmatic tree model of hematopoiesis is increasingly recognized to be limited as it is based on heterogeneous populations and largely inferred from non-homeostatic cell fate assays. Here, we combine persistent labeling with time-series single-cell RNA-Seq to build the first real-time, quantitative model of *in vivo* tissue dynamics for any mammalian organ. We couple cascading single-cell expression patterns with dynamic changes in differentiation and growth speeds. The resulting explicit linkage between single cell molecular states and cellular behavior reveals widely varying self-renewal and differentiation properties across distinct lineages. Transplanted stem cells show strong acceleration of neutrophil differentiation, illustrating how the new model can quantify the impact of perturbations. Our reconstruction of dynamic behavior from snapshot measurements is akin to how a Kinetoscope allows sequential images to merge into a movie. We posit that this approach is broadly applicable to empower single cell genomics to reveal important tissue scale dynamics information.

**Highlights:**

- Cell flux analysis reveals high-resolution kinetics of native bone marrow hematopoiesis
- Quantitative model simulates cell behavior in real-time and connects it with gene expression patterns
- Distinct lineage-affiliated progenitors have unique self-renewal and differentiation properties
- Transplanted HSCs display accelerated stage- and lineage-specific differentiation

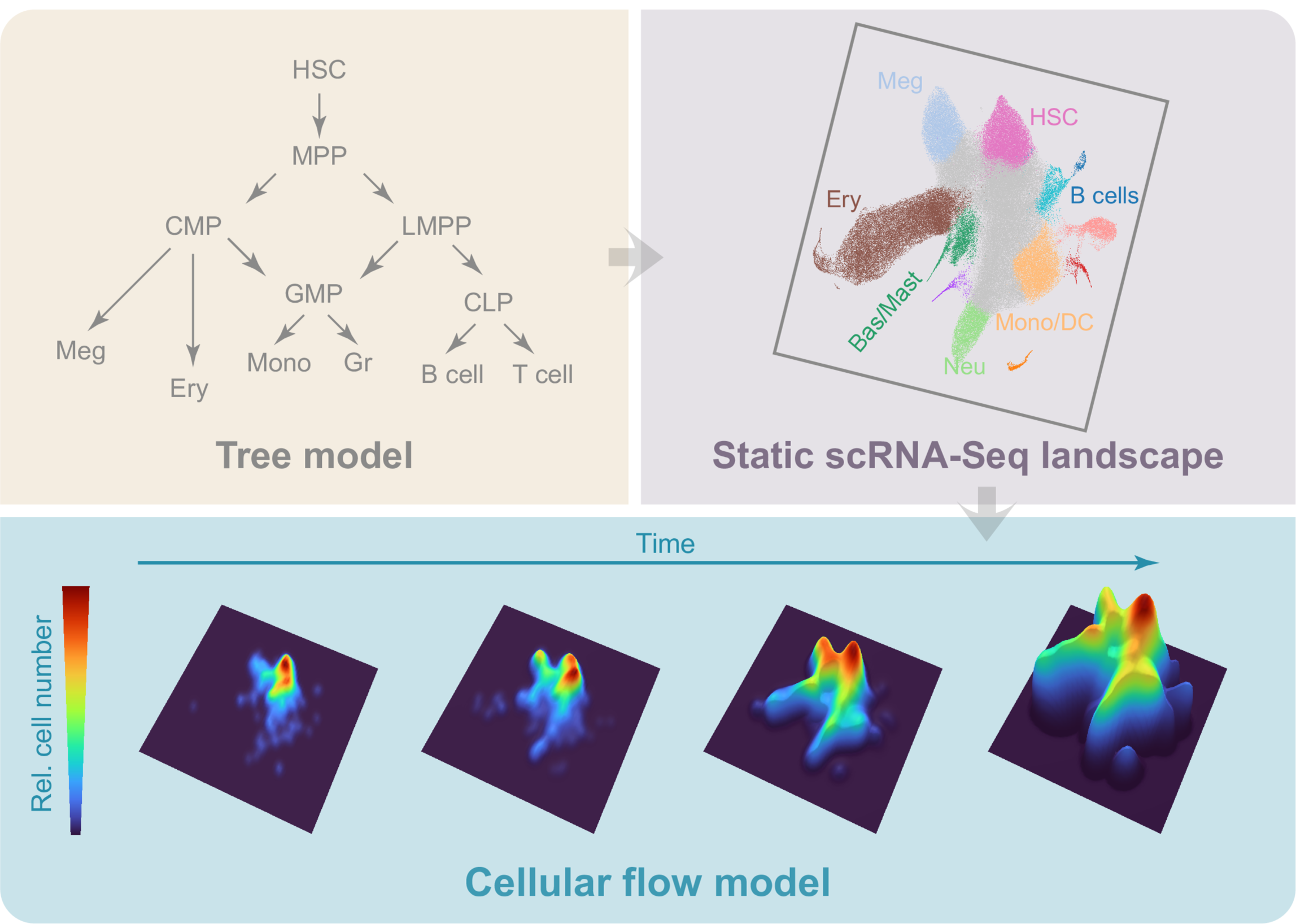

## Introduction

A continuous flow of cells replenishes blood cells throughout life to maintain homeostasis. This flow originates from the hematopoietic stem cells (HSCs) and progresses through a complex hierarchy of multipotent, bipotent and unipotent progenitors, together called hematopoietic stem and progenitor cells (HSPCs). Decades of research have allowed to immunophenotypically identify HSPCs and define their functionality, thus positioning them within the hematopoietic hierarchy and establishing the ’hematopoietic tree model’ (Eaves, 2015; Notta et al., 2016).

The hematopoietic tree, while undeniably useful, is a static and qualitative model of a highly dynamic process. Previous work (Busch et al., 2015) paved the way to real-time modelling of HSPC dynamics under native conditions. The study induced a persistent fluorescent reporter gene within the HSC compartment and followed label propagation into downstream progenitor and mature cells with flow cytometry. However, immunophenotyping has limited resolution, and HSPC populations defined by flow cytometry are known to be functionally heterogeneous. This is particularly evident within common myeloid progenitors (CMP) (Paul et al., 2015; Perié et al., 2015) and lymphoid-primed multipotent progenitors (LMPP) (Klein et al., 2022; Nestorowa et al., 2016) as revealed by scRNA-Seq and transplantation experiments. Further high-throughput scRNA-Seq studies charted putative gradual molecular transitions from HSCs toward 8 distinct lineages (Dahlin et al., 2018) including specific stages of erythroid differentiation (Tusi et al., 2018). More recently, lineage tracing and scRNA-Seq were combined to show that molecular states captured by scRNA-Seq are predictive of progenitor fate potential when assessed *in vitro* (Wang et al., 2022; Weinreb et al., 2020; Yeo et al., 2021), but gaining insight into single-cell fates *in vivo* during homeostasis is more challenging (Pei et al., 2020).

While scRNA-Seq offers high-resolution, it is typically used to obtain snapshot measurements lacking temporal information. Here, we combined scRNA-Seq with an inducible HSC-labelling system allowing label-propagation analysis of the downstream progeny during steady-state hematopoiesis. We measured the real-time dynamics of label accumulation across the stem and progenitor cell landscape and built cellular flow models capturing self-renewal and differentiation rates. We find that cell output is maintained *via* lineage-specific mechanisms. By taking advantage of the available molecular information, we also construct continuous models to associate the gene expression changes with cell behaviors such as increased proliferation or accelerated differentiation, thus directly connecting tissue and cellular behavior with the underpinning layer of molecular processes. Finally, we demonstrate that our reference model, unlike immunophenotypic data, is transferable and applicable to different datasets. To showcase this, we analyze transplanted stem cell progeny and pinpoint drastic upregulation of differentiation rates in specific lineages.

## Results

### *Hoxb5-Cre^ERT2^-Tomato* reporter tracks HSC differentiation over time

To analyze the HSPC dynamics, we employed a heritable fluorescent label approach (based on principles from (Busch et al., 2015)), in which an inducible HSC-specific CRE excises a STOP cassette in *Rosa26-LoxP-STOP-LoxP-tdTomato* (*R26^LSL-tdTomato^*) reporter to permanently label HSCs and the label expression is subsequently inherited by their downstream progeny. To achieve this, we generated the *Hoxb5^CreERT2^* mouse allele where the CRE-ERT2 protein is expressed from the HSC-specific *Hoxb5* gene (Figure 1A), following a similar strategy previously employed to express mCherry from the *Hoxb5* locus without affecting *Hoxb5* expression (Chen et al., 2016).

**Figure 1.**
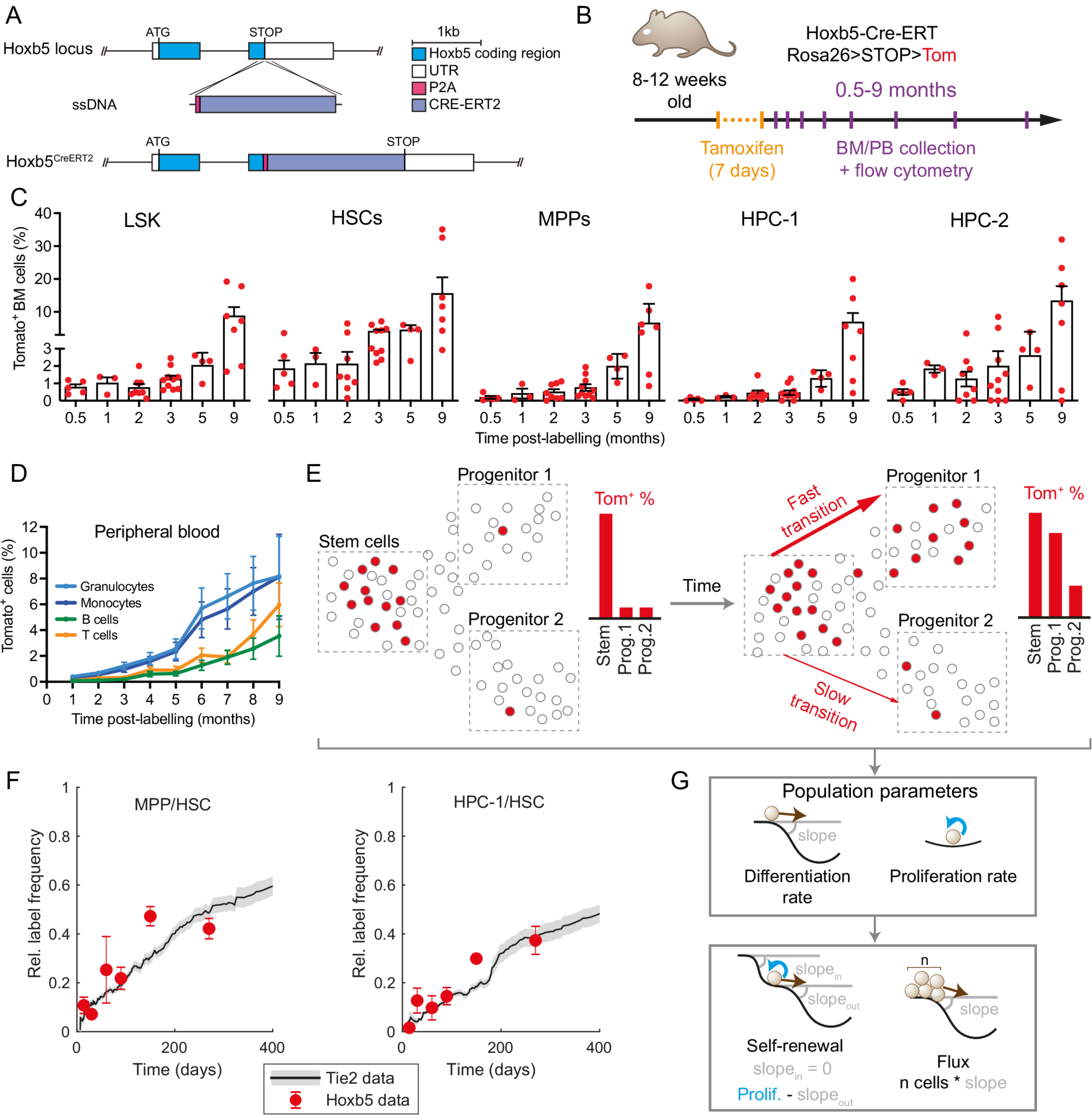
Hoxb5-Tom persistent labelling system enables time-resolved tracking of stem cells and their progeny. (A) Diagram of the genetic construct used to introduce the inducible and persistent Hoxb5-Tom label in the respective mouse line. (B) Schematic of the time-course experiment analyzing Hoxb5-Tom label frequency in the indicated populations of mouse bone marrow (BM) and peripheral blood (PB). Upon tamoxifen administration, Hoxb5-expressing cells are labelled with heritable Tom expression. (C) Fractions of Tom^+^ cells in the HSPC subpopulations within the bone marrow at indicated time-points after label induction. Mice were analyzed at 0.5 (n=5), 1 (n=3), 2 (n=8), 3 (n=10), 5 (n=4) and 9 (n=7) months after label induction. Dots represent individual mice and bars indicate mean ± SEM. (D) Fractions of Tom^+^ cells in peripheral blood analyzed at the indicated time-points after label induction. Shown as mean with error bars denoting SEM of 4-32 animals. (E) Diagram portraying the concept of inferring population dynamics from heritable label propagation. The rate of label accumulation in the downstream compartments is proportional to differentiation rate between the compartments. (F) Comparison of Tie2-YFP and Hoxb5-Tom label progression displayed as relative labelling frequency between MPP or HPC-1 and HSC compartments. Red dots - Hoxb5-Tom data points, grey line - rolling average for matching Tie2-YFP data, as published previously (Barile et al., 2020). (G) Diagrams portraying key population parameters together with a geometric interpretation in context of the Waddington landscape.

We next combined the *Hoxb5^CreERT2^* allele with the *R26^LSL-tdTomato^*reporter (Madisen et al., 2010) to generate *Hoxb5^CreERT2^*; *R26^LSL-tdTomato^* mice (for simplicity referred to as Hoxb5-Tom mice), which allow for inducible labelling of HSCs *in situ* at a specific time-point by tamoxifen administration and tracking HSC progeny over time (Figure 1B-D). To validate this system, we treated Hoxb5-Tom mice with tamoxifen and monitored tdTomato (Tom) expression in HSCs and their subsequent progeny within the HSPC sub-populations in the bone marrow (BM) and differentiated cell types in the peripheral blood (PB) at indicated intervals (Figure 1B-D, S1A-C). Upon tamoxifen administration, we observed a specific labelling of the HSC compartment (with the frequency of 1.8%), which over 2 months gradually accumulated in downstream cell compartments (Figure 1C-D). Labelled differentiated cells are detectable in PB within 1-2 months after labeling HSCs; with granulocytes and monocytes being the first emerging populations and T and B cells appearing later. We observed stable labelling for at least 9 months after the treatment (Figure 1C-D, S1A-B), indicating that the label is persistent and inert.

Computational inference of population dynamics relies on a simple principle (Figure 1E): as heritable label propagates down from the label-rich upstream compartment it follows differentiation, thus rapid transitions cause fast label equilibration and vice versa (see methods). To benchmark our new model, we compared flow cytometry data obtained from tamoxifen-treated Hoxb5-Tom mice with previously published results (Busch et al., 2015) of analogous label propagation obtained with the Tie2-YFP mouse line. As shown in Figure 1F, our data are highly consistent for both MPP/HSC and HPC-1/HSC relative abundances across the entire time range. Altogether, we established a new mouse system for inducible, persistent labelling of HSC progeny in the BM, thus unlocking our next goal - modelling of population dynamics.

### A unified reference HSPC landscape with time-resolved differentiation

Having validated the HoxB5-Tom system, we designed a strategy to capture scRNA-Seq profiles of cells traversing the HSPC landscape in real-time (Figure 2A). We harvested BM from tamoxifen-treated mice at 9 time-points ranging between 3 days (providing just enough time for Tom expression) and 269 days, when the label is mostly equilibrated. At each time-point we sorted Lin^-^ cKit^+^ or Lin^-^Sca1^+^ sub-population from the bone marrow, which contains all stem cells and a broad view progenitor cells (Dahlin et al., 2018) (Figure S1D). To ensure accuracy and reproducibility, we profiled multiple independent biological replicates for each time-point (36 animals in total). While our focus was labelled Tom^+^ cells, we also profiled Tom^-^ cells at each time-point to obtain the background cell density. We generated a common reference landscape by integrating all single-cell profiles followed by clustering and embedding in a UMAP projection (Figure S2F). Clusters disjointed from the main landscape body (mostly mature cell types) and those representing technical artifacts (e.g. doublets or dying cells) were excluded. The refined landscape (>115,000 cells) served as a basis of our analysis (Figure 2B, unfiltered data in Figure S2F,G).

**Figure 2.**
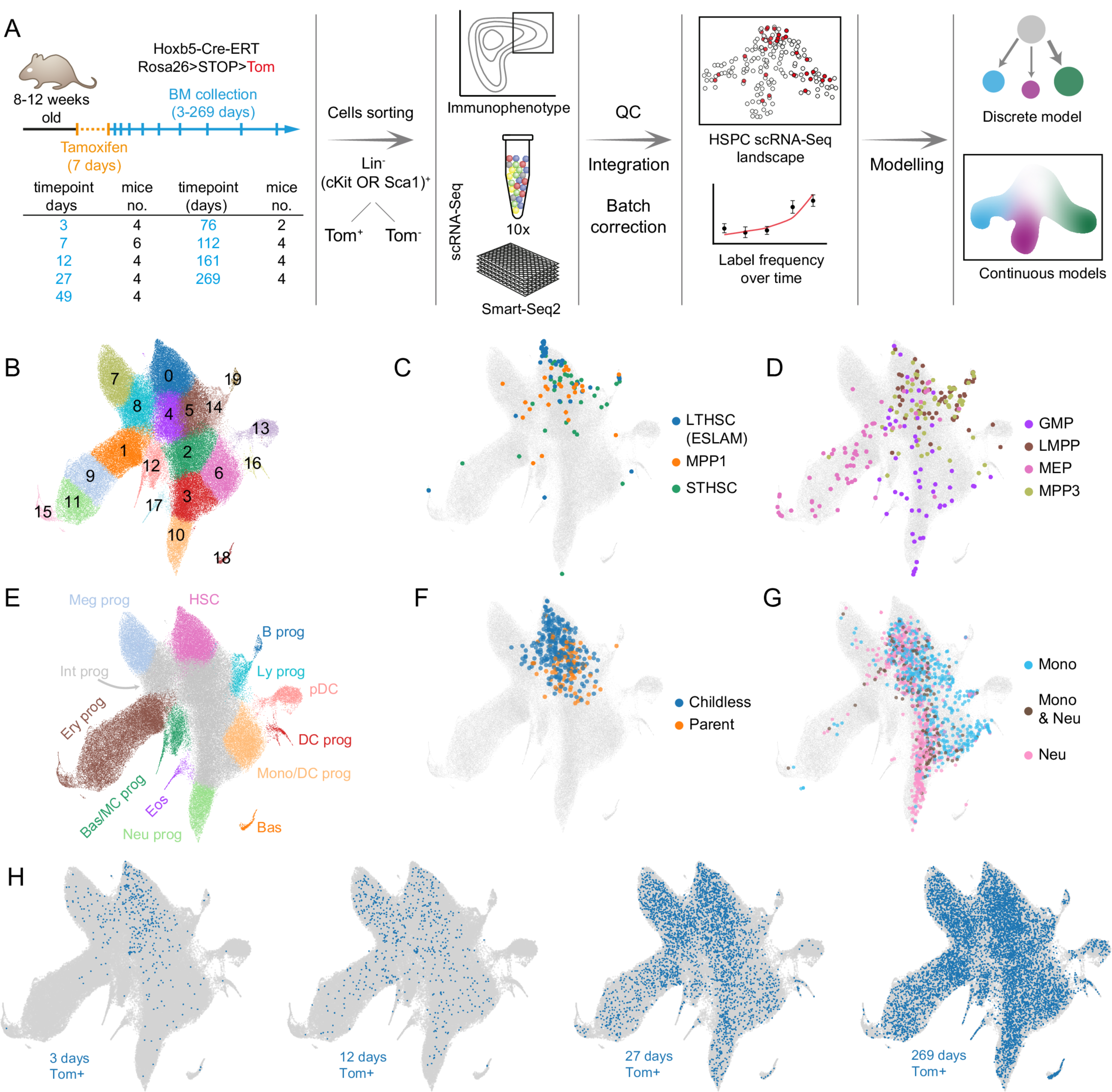
Time-resolved reference HSPC landscape at single-cell level. (A) Experimental design for HSPC dynamics analysis with flow cytometry and scRNA-Seq. Table indicates specific time-point and the number of mice (replicates) used Tom^+^ scRNA-Seq analysis, 2 mice in each time-point were used for the Tom^-^ fraction estimation. (B) UMAP projection of the integrated HSPC scRNA-Seq landscape (all Tom^+^ and Tom^-^ cells combined) with color-coded clusters. Outlier or aberrant clusters were removed for clarity (see Figure S2F,G). (C,D) Projection from B in grey, with embedded and color-coded immunophenotypic sub-populations from (Nestorowa et al., 2016) data. Up to 60 cells in each category are plotted. All cells are plotted in the Figure S4A. (E) Manual annotation of the landscape in B. Most differentiated clusters with clearly defined lineage markers are color-coded, intermediate undifferentiated states are shown in grey (Int prog), cluster containing HSCs is shown in pink. (F) Projection from B in grey, with embedded and color-coded HSCs with no detected cellular output (Childless) or contributing to haematopoiesis (Parent) following 5-FU challenge in mice (data from (Bowling et al., 2020)) (G) Projection from B in grey, with embedded and color-coded cKit^+^ progenitors, based on their output in lineage tracing *in vitro* cultures. Color-coded points correspond to cells harvested at day 2 with sufficient clonal information available at day 4 and day 6 of culture. Data from (Weinreb et al., 2020). (H) Projection from B in grey, with Hoxb5-Tom^+^ cells harvested at indicated time-points shown in blue. Abbreviations: B prog - B cell progenitor, Bas - basophils, Bas/MC prog - Basophil and Mast Cell progenitors, DC prog - dendritic cell progenitors, Eos - eosinophils, Ery prog - erythroid progenitors, HSC - hematopoietic stem cells, Int prog - intermediate progenitors, Ly prog - lymphoid progenitors, Meg prog - megakaryocyte progenitors, Mono/DC prog - monocyte and dendritic cells progenitors, Neu prog - neutrophil progenitors, pDC - plasmacytoid dendritic cells

To place our data within the broader scope of hematopoietic research and extend its interpretability, we provide multiple layers of annotation. The manual annotation (Dahlin et al., 2018; Weinreb et al., 2020) uses lineage marker expression (Figure S2A,B) and HSC-score (molecular signature of long-term repopulating HSCs (Hamey and Göttgens, 2019)) to highlight the upstream cluster containing HSCs (Figure S2C) (cluster 0) and 8 terminal clusters (Figure 2E), where clear expression of definitive markers is observed (Supplementary Table 1). To add more functional information, we mapped external scRNA-Seq datasets using our Cellproject package. Firstly, we overlaid canonical immunophenotypic annotation (data from (Nestorowa et al., 2016)) comprising: highly purified LT-HSCs, multipotent progenitors (MPPs) 1 and 3, ST-HSCs, granulocyte-monocyte progenitors (GMPs), LMPPs and megakaryocyte-erythroid progenitors (MEPs) (Figure 2C,D, S4A,B). Secondly, we highlighted cell states associated with specific cell fate outcomes based on *in vitro* lineage tracing experiments (Weinreb et al., 2020) (Figures 2G and S4C). Importantly, the *in vitro* cell potency is broadly aligned with the manual cluster annotation. Finally, we included information about the active/inactive HSC status under proliferative challenge based on lineage tracing data from (Bowling et al., 2020) (Figure 2F). Together, these annotations place cell clusters in the functional context, essential for interpretation of the population dynamics models discussed below.

The HSPC landscape split by time-point shows clear propagation of labelled cells (Figure 2H, quantification for all time-points is shown in Figure S3B), in agreement with the theory (Figure 1E). Certain areas (e.g. clusters 8 and 7) very quickly accumulate labelled cells, others are slower (clusters 11 or 10) and some very slow (clusters 13 or 14). Eventually the label largely equilibrated, as compared to the Tom^-^ population (Figure S3A). Importantly, cell populations defined in this way are much more molecularly homogeneous, in contrast to conventional flow cytometry gates (Figure S4A,B) (Nestorowa et al., 2016; Paul et al., 2015; Rodriguez-Fraticelli et al., 2018). To provide a quantitative description of population dynamics, we employed two types of models: discrete and continuous, each built for specific purpose. The former captures dynamics across the entire compartment and intuitively combines the old models of hematopoiesis with a new quantitative view. It also serves as a necessary reference for the latter, a more advanced continuous modelling approach, which focuses on specific trajectories, but provides cellular flux parameter estimates for each single cell and thus directly connects single cell transcriptomic profiles with tissue-scale cellular behavior.

### Discrete model reveals HSPCs with lineage-specific patterns of self-renewal and differentiation

To capture flow of cells through HSPC compartment in real time, we utilized the label propagation principles to build a discrete model consisting of multiple, interconnected cell clusters (Figure 3A- C). We explain two variables changing over time: initial labelling frequency (Tom^+^ cells) and size (Tom^-^ cells) per cluster. The model considers two basic properties of each cluster: *net proliferation* (difference between proliferation and death) and *differentiation rates* (ingoing and outgoing transitions among clusters). Additionally, we introduce two derived parameters that are useful for interpreting cell behavior (Figure 1D). *Residence time*, a metric of self-renewal, describes the time required for the cluster to shrink by 63% (to 1/e of original size, where e is the Euler’s number) in absence of any inputs. Finally, *flux* depicts the number of cells transported in a unit of time. We limited the number of differentiation parameters by assuming that cells travel only between adjacent clusters (i.e. with highest PAGA connectivities – Figure 3A, (Wolf et al., 2019)).

**Figure 3.**
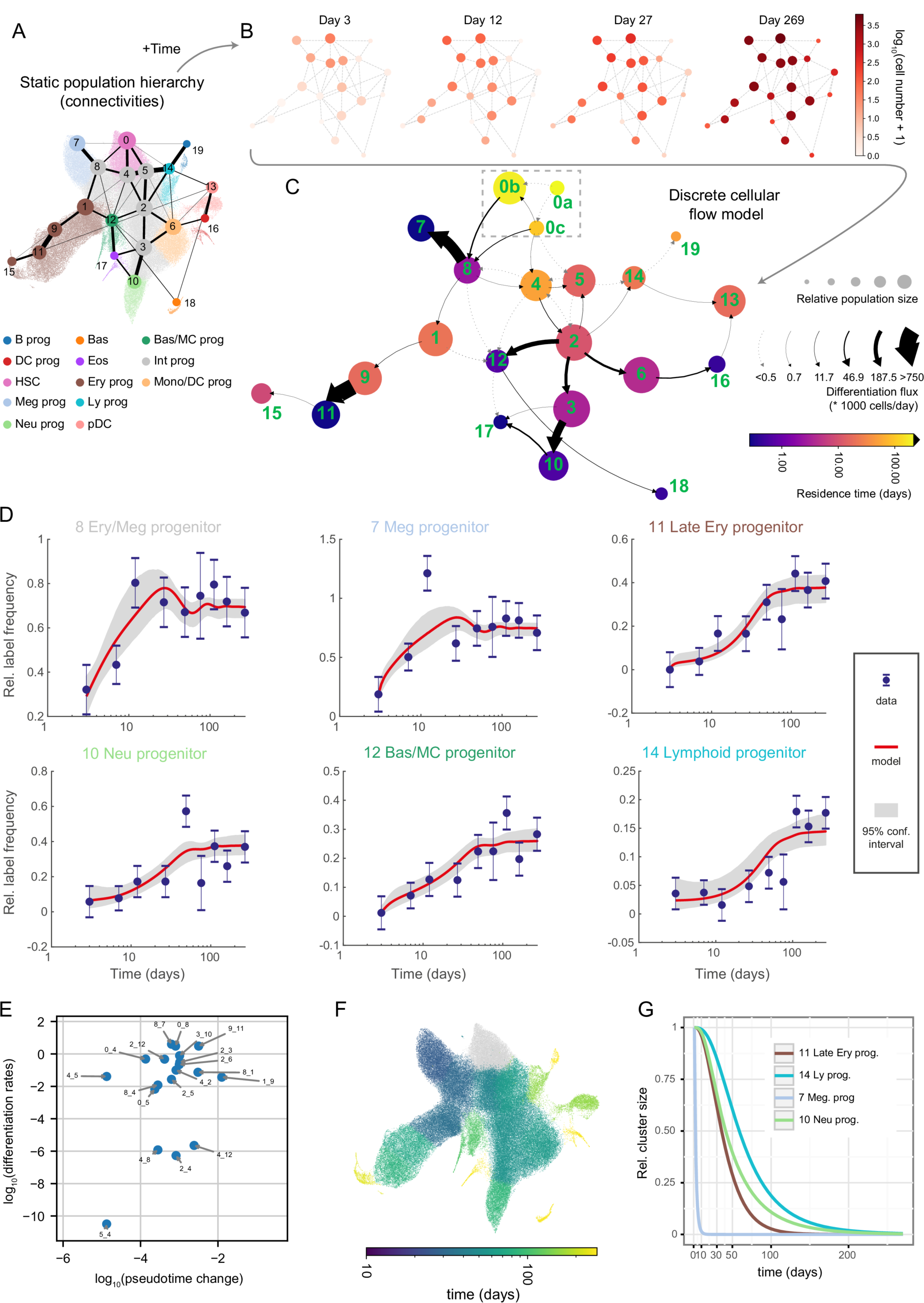
Quantitative discrete model of the HSPCs highlights progenitor-specific self- renewal and differentiation properties. (A) Annotated UMAP projection overlaid with PAGA graph abstraction view of the HSPC landscape. The graph shows putative transitions between clusters (related to Figure 2B). (B) The absolute number of labelled cells observed in each cluster over time displayed as a graph view from A. 4 out of 9 time-points are shown for clarity. (C) Graph abstraction view of the discrete cellular flow model. Size of the nodes is proportional to square roots of relative cluster size, node color is proportional to the residence time (log-scale), arrows indicate differentiation directions, arrow stem thickness is proportional to cell flux. Note: cluster 0a is fully self-renewing and thus exhibits infinite residence time. (D) Best discrete model fit (with 95% confidence intervals) for relative Tom^+^ label frequency in chosen clusters. Error bars indicate pooled standard error of the mean. (E) Scatter plot showing relation of pseudotime distance to differentiation rates. Only clusters 0-12 and differentiation rates greater than 10^-12^ are shown. Please note that in case of the transitions between clusters 4 and 8 two differentiation rates are plotted (each direction). (F) UMAP projection of the HSPC landscape, with cells color-coded by simulated time required for 1 cell to reach corresponding cluster starting from cluster 0. Please mind that the color is logarithm-scaled. (G) Simulated relative cluster size of chosen clusters following ablation of cluster 0.

Of note, we observed changes in relative cluster size over-time, in particular a quick increase (>50% in <20 days) in clusters 0, 7 and 8 and a coordinated decrease in other major clusters (Figure S6), (Sánchez-Aguilera et al., 2014). Previous tamoxifen-based label propagation studies coupling also observed a quick rise in ST-HSC, MPP2 and MPP3 total numbers (Figure S6B), but no explanations were provided (Barile et al., 2020). Consistent with recovery from cell depletion caused by tamoxifen interference with JAK/STAT signaling (Sánchez-Aguilera et al., 2014), this pathway was most active in the depleted clusters 7, 8 in addition to cluster 0 (Figure S7H). To assess how recovery from cell depletion may influence model parameters, we compared our main model with a bi-phasic fit which permits a switch in differentiation/proliferation rates between the recovery phase and homeostasis (after 27 days), which shows that 14 out of 58 rates change (Figure S7D-G). We thus explain and account for a previously overlooked side-effect of using tamoxifen for label induction.

We formulated our main model into a graph in Figures 3C and S7A, where node sizes are proportional to the average cluster size, node color indicates self-renewal (or net proliferation in Figure S7A) and arrows indicate cell flux (differentiation rate in Figure S7A). Interestingly, differentiation rates do not correlate with similarities between gene expression states (Figures 3E, S7B), indicating that inferring real-time dynamics requires temporal information.

Let us first consider cluster 0 as a single unit (grey box, Figure 3C), whose substructure we will discuss in the next section. The compartment-wide view clearly shows lineage-specific dynamics (Figure 3C). The definitive megakaryocyte progenitors emerge through a rapid transition via the fast-proliferating cluster 8, which also more slowly generates erythroid cells (cluster 1). The erythroid lineage is maintained by including additional stages with considerable self-renewal (clusters 1 and 9) and proliferation (cluster 9), followed by fast differentiation between clusters 9 and 11. The myeloid progenitors flow from cluster 0 either into cluster 4 or via a shared route with the erythroid and megakaryocytic progenitors in cluster 8, with gradually increasing differentiation rates from cluster 2 onward. Thus, the myeloid branch similarly to the erythroid one, just like the erythroid one, employs additional progenitor populations, albeit with lower proliferation rates (Figure S7A).

By contrast the lymphoid trajectory shows low proliferation and is limited by slow transitions via clusters 5 and 2 into cluster 14 (which overlaps mostly with a subset of MPP4 cells). Similarly slow is the plasmacytoid dendritic cell (cluster 13, pDCs) differentiation through the lymphoid cluster 14 and myeloid clusters 6 and 16. The emergence of mast cell, basophil and eosinophil progenitors in the adult bone marrow is unclear (Hamey et al., 2021; Wu et al., 2022). Our results reveal that basophil and mast cell progenitors (cluster 12) are continuously generated and originate mostly by a transition from cluster 2. Furthermore, despite limited cell numbers, we observed some label accumulation in eosinophils (cluster 17), primarily originating from neutrophil progenitors (cluster 10).

Interestingly, residence time (self-renewal) varies widely across the HSPC landscape, with lineage-specific patterns (Figure 3C). As expected, cluster 0 contains the only perfectly self-sustaining population; intermediate populations show an extensive range of residence times, from just 2.5 days for Erythroid/Megakaryocytic progenitor (cluster 8), 11 days for Monocyte/Granulocyte progenitors (cluster 2) and up to 53 days for the medial cluster 4. The latter example falls close to the residence time previously estimated for MPPs (70 days)(Busch et al., 2015) and highlights that progenitors can also show considerable self-renewal. Importantly, cells in cluster 8, 2 and 4 largely fall within the immunophenotypic CMP and MPP definitions (Figures 2C-D and S4A,B), indicating that these gates capture populations with vastly different dynamics. We also note that among some intermediate clusters our model permits a degree of forward and backward differentiation suggesting a balance (or oscillations) between the states. To conclude, diverse hematopoietic progenitors exhibit widely different, lineage-dependent dynamics illustrating the distinct mechanisms maintaining cell output.

### The top compartment composition changes over time

The top cluster 0, based on the immunophenotype annotation (Figure 2C), contains virtually all LT-HSC and a large subset of ST-HSC and MPP1 cells. The overall cluster size increases over time (Figure S5C), following the expansion of ST-HSCs and MPP3s (Figure S6B) (Barile et al., 2020). Surprisingly, the Hoxb5-Tom labelled cells within cluster 0 grow almost exponentially (Figure S5B), which mirrors the behavior of Tie2-YFP labelled LT-HSCs (Barile et al., 2020) and is consistent with the observation of dramatic expansion of Hoxb5-, Tie2- or Fgd5-labelled cells in ageing animals (Zhang et al., 2020). This suggests that Hoxb5 and Tie2 systems mark, in addition to the canonically quiescent LT-HSCs, a subset of stem cells with high self-renewal or proliferation capacity.

To provide insight into cluster 0 sub-structure, we tested multiple models and put forward a potential explanation, which assumes a logistic growth for cluster 0 and three sub-clusters within in it: a top, perfectly self-renewing cluster 0a, megakaryocyte & myeloid-biased cluster 0b and multipotent 0c. We constrained cluster 0a size and differentiation rate to match previously reported LT-HSC numbers but left clusters 0b and 0c sizes unconstrained. Cluster 0c remains stable over time but it proliferates quickly and feeds both downstream progenitors and cluster 0b, which in turn grows over time (Figure S5D). Hence, the flux between clusters 0b and 8 increases with mouse age. This is in line with the increased myeloid output (Benz et al., 2012; Muller-Sieburg et al., 2004) and relative proportion of megakaryocyte-biased and myeloid-biased HSCs in aged animals (Yamamoto et al., 2018). Of note, cluster 0b shows high self-renewal (residence time of 180 days), consistent with high repopulation potential of lineage-biased HSCs (Yamamoto et al., 2018). Altogether, our discrete model in addition to faithful recapitulation of cell flux through the HSPC compartment also provides a possible explanation of ageing-associated changes in HSC behavior.

### Continuous model of hematopoiesis connects dynamics of gene expression with cell behavior

Unlike immunophenotyping, scRNA-Seq contains transcriptome-wide profiles for each cell. While the discrete model provides compartment-wide dynamics, its utility for gene expression analysis is limited. To address this issue, we employed a continuous model based on the Pseudodynamics framework (Fischer et al., 2019). For tractability, we considered one lineage at a time, based on cells with highest fate probabilities towards each lineage (Lange et al., 2022; Setty et al., 2019)(Figure 4A,B, S8A,B, S11).

**Figure 4.**
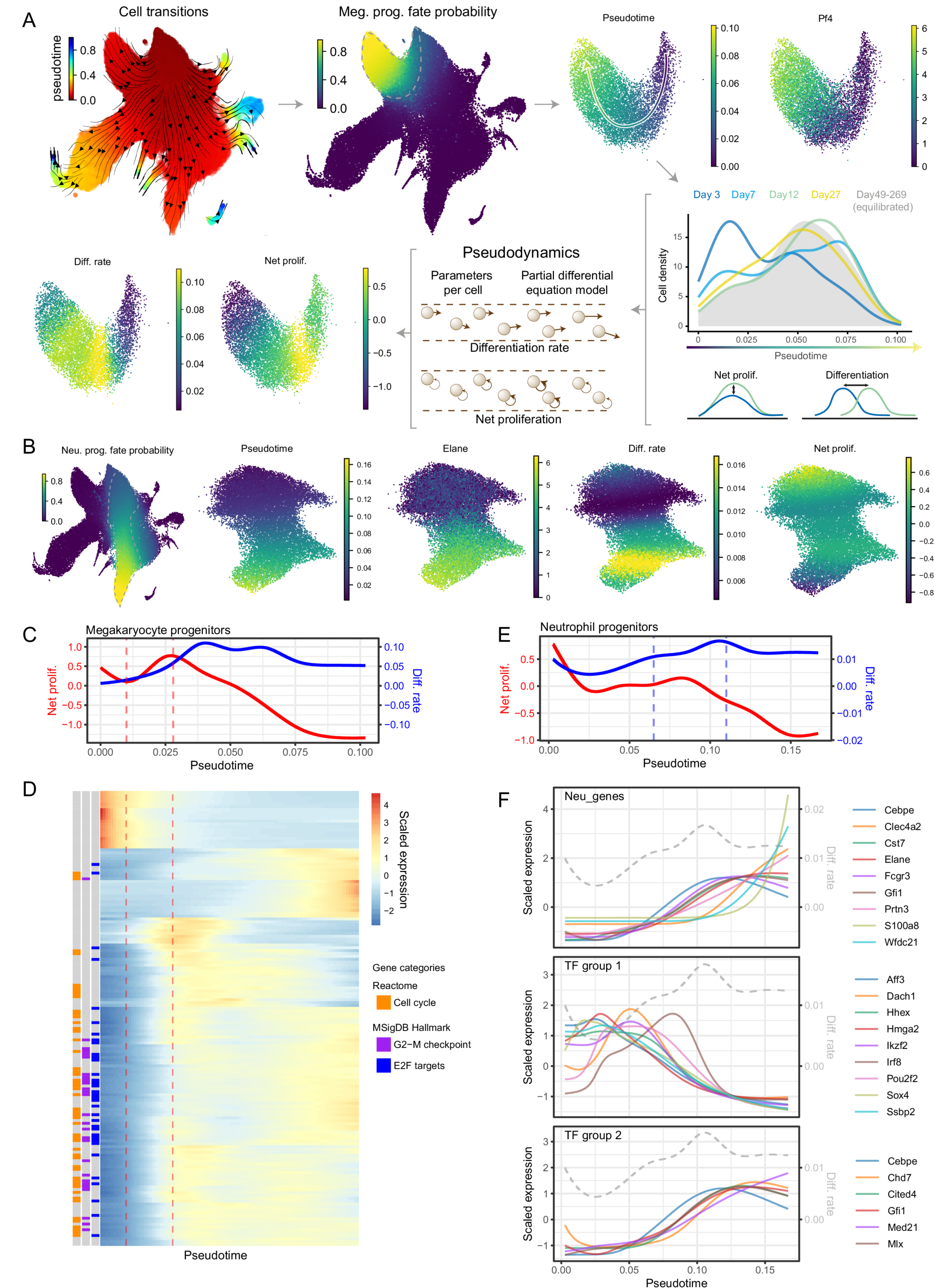
Continuous models capture single cell growth and differentiation rates alongside their molecular state. (A) Diagramatic representation of megakaryocyte trajectory analysis with pseudodynamics. Following the arrows: putative cell transitions (pseudotime kernel) were used to estimate megakaryocyte cell fate, from which megakaryocyte trajectory was isolated (dashed line). Along the pseudotime cell densities were computed for each time-point (color-coded density profiles) and analyzed using the pseudodynamics framework providing differentiation and net proliferation rate estimates for each cell. (B) (left) UMAP projection of the HSPC landscape color- coded by cell fate probability of neutrophil lineage (estimated with pseudotime kernel, see A). Panels on the right show UMAP projections of isolated neutrophil trajectory color-coded by indicated parameters or gene expression. (C) Pseudodynamics fitted net proliferation parameter (red) and differentiation rate parameters (blue) along pseudotime for megakaryocyte trajectory. Vertical lines indicate the region of interest. (D) Heatmap of genes differentially expressed around the region of interest shown in C. Left columns indicate genes belonging to enriched gene categories - E2F target (FDR <10^-38^), G2-M checkpoint (FDR <10^-24^) and cell cycle (FDR <10^-38^). (E) Pseudodynamics fitted net proliferation (red) and differentiation rate (blue) parameters along pseudotime for neutrophil trajectory. Vertical lines indicate the region of interest. (F) Fitted gene expression values along pseudotime for neutrophil markers and two TF groups shown in (full analysis in Figure S9A). Grey, dashed line indicated differentiation rates shown in E. Gene expression was scaled around the mean.

The continuous model assigns differentiation and net proliferation rates to each cell (Figure 4A) by solving partial differential equations describing cell densities along pseudotime over real-time. Hence, model parameters and gene expression share a common pseudotime (and real-time) axis, enabling direct comparison. Of particular interest are states (i.e. pseudotime ranges) with changes in proliferation or differentiation rates. An increase in proliferation rates indicates an expansion stage, whereas a rise in differentiation rates marks a potentially irreversible molecular transition.

We set out to analyze gene expression dynamics occurring at such changes in cell behavior. For brevity we focus on the megakaryocyte and neutrophil trajectories but also provide analogous analyses for erythroid and monocytic/dendritic lineages (Figures S8,S9). As shown in Figure 4A,C megakaryocyte progenitors show characteristic changes in growth and differentiation rates. Cells rapidly increase their net proliferation early on, ahead of the peak in differentiation and around the stage where *Pf4* (megakaryocyte marker) mRNA becomes detectable. In this growth phase, we identified 170 dynamically expressed genes (similar analysis of the differentiation phase in Figure S8C-D) with distinct patterns along pseudotime (Figure 4C-D). These genes are strongly enriched for cell growth and proliferation genes with almost all of them showing an upward trend in the relevant pseudotime range. This serves as a proof of principle, as the model based solely on total cell numbers, predicts the growth stage matching the respective gene signature.

While following the neutrophil differentiation kinetics (Figure 4B,E), we found gradually increasing differentiation rates (blue line) accompanied by a complex pattern of gene expression. Indeed, we observed two phases of neutrophil-affiliated gene expression (Figure 4F), with *Cebpe*, *Cst7*, *Elane*, *Fcgr3*, and *Gfi1* appearing almost simultaneously at the onset of differentiation, while *Clec4a2*, *Wfdc21*, *S100a8* increasing at different intervals later. To gain insight into potential mechanisms regulating the process, we scrutinized transcription factors with dynamic expression along the trajectory (Figure S9A) and classified them into 4 groups based on their distinct expression patterns. Group 2 (Figure 4F) largely mirrored the expression of early neutrophil markers described above, and reassuringly contained the *Gfi1* factor, a key determinant of the neutrophil fate (Olsson et al., 2016). Group 3 (Figure S9B) contained factors with the highest expression in the most immature HSPCs (e.g. *Gata2*, *Hlf*, *Meis1*) and showed early and nearly synchronous decay in expression, suggesting involvement in self-renewal. Finally, Group 1 (Figure 4F) TFs exhibit unique patterns of expression with peaks at different stages, all of which ultimately decaying as late neutrophil markers appear. These contain multiple TFs associated with specific lineages such as: *Irf8* (Monocyte/DC fate (Olsson et al., 2016)), *Aff3* (lymphoid/B cells (Ma and Staudt, 1996)), *Dach1* (myeloid (Amann-Zalcenstein et al., 2020)), *Hmga2* (myeloid, erythroid, megakaryocytic (Kumar et al., 2019), *Pou2f2* (lymphoid/B cells (Novershtern et al., 2011) or are important for HSPC self-renewal, including *Ikzf2* (Park et al., 2019) or *Ssbp2* (Li et al., 2014). Thus, our analysis indicates that progenitors exhibit transient expression of major lineage determinants at specific differentiation stages on their way to becoming neutrophils (see *Gfi1*, *Flt3*, *Irf8* in Figure S9D,E). Early accumulation of these factors is correlated with increased differentiation rate but eventually a single programme takes over and accelerates the differentiation even further. To conclude, the continuous model unlocks the access to full single cell transcriptome data, and thus enables integrated analysis of cellular and molecular dynamics, revealing new mechanistic insights into cell behavior during differentiation.

### HSPC models simulate cell journeys in real-time consistent with basic properties of hematopoiesis

Mathematical models combined with our new datasets offer unique prediction capabilities allowing us to unravel fundamental facets of hematopoiesis. Specifically, we focused on computing cell journeys in real-time and consequences of cluster ablation. Firstly, we estimate the ’average journey times’ with the discrete model. We placed a single cell in cluster 0 and computed the average time required to accumulate one cell for each target cluster. As shown in Figures 3F and S7C, time required to reach terminal population varies widely between lineages. For instance, reaching Meg progenitors (cluster 7) requires 27 days, neutrophil progenitors (cluster 10) or late erythroid progenitors (cluster 11) >80 days and finally producing pDCs takes about 150 days. Secondly, we predict what would happen if, under normal conditions, the self-renewing cluster 0 was ablated. As expected, without cluster 0 input, downstream cluster sizes would gradually decline over time (Figure 3G), due to limited self-renewal of intermediate progenitors. As we described above, progenitor self-renewal is lineage-specific, hence corresponding clusters wane at different rates, with megakaryocyte progenitors depleted to 50% after 2-3 days, whereas lymphoid progenitors are maintained for >50 days.

Altogether, the hierarchy revealed by our model is consistent with current hematopoiesis research (Eaves, 2015; Rodriguez-Fraticelli et al., 2018; Tusi et al., 2018; Weinreb et al., 2020). Furthermore our model estimates lead to predictions that agree with basic properties of hematopoiesis inferred from transplantation (Notta et al., 2016; Rodriguez-Fraticelli et al., 2018) or cell culture (Notta et al., 2016; Weinreb et al., 2020) experiments such as the order of lineage emergence. The time-frame of the process is expectedly much longer but is compatible with previous studies of HSPC dynamics in the native context (Busch et al., 2015). This important validation demonstrates that our completely new approach is anchored firmly in the long tradition of hematopoiesis research, yet at the same time produces profound new insights, and unlocks previously impossible means of performing truly quantitative comparisons.

### Integrative model resolves effects of transplantation HSPC dynamics

Our models serve as a reference framework for native hematopoiesis, uniquely capable of transferring information across experiments and systems. To demonstrate this utility, we analyzed previously published data (Dong et al., 2020) using scRNA-Seq to track the progeny of highly-purified HSCs in transplanted animals over time (Figure 5A). After integrating the scRNA-Seq profiles into our reference landscape (Figure 5B-F), we derived cell frequencies per cluster at day 3, and used the discrete model to predict the cell abundance expected under ’normal’ conditions (Figures 5 and S10). While some general features match normal hematopoiesis, for instance megakaryocyte progenitors being the first emerging lineage, it is clear that, under transplantation conditions, cells differentiate much faster in most directions, particularly towards the neutrophil fate (Figure 5G, cluster 10). The erythroid lineage behaves differently; while early megakaryocyte and erythrocyte differentiation is accelerated upon transplantation (Figure 5G, cluster 8), late erythroid progenitor cell differentiation is stalled, compared to the steady-state counterparts (Figure 5G, cluster 11). In conclusion, we demonstrated that our model can be easily applied to other datasets, and provide quantitative predictions and interpretation, which would not be otherwise available from static measurements alone.

**Figure 5.**
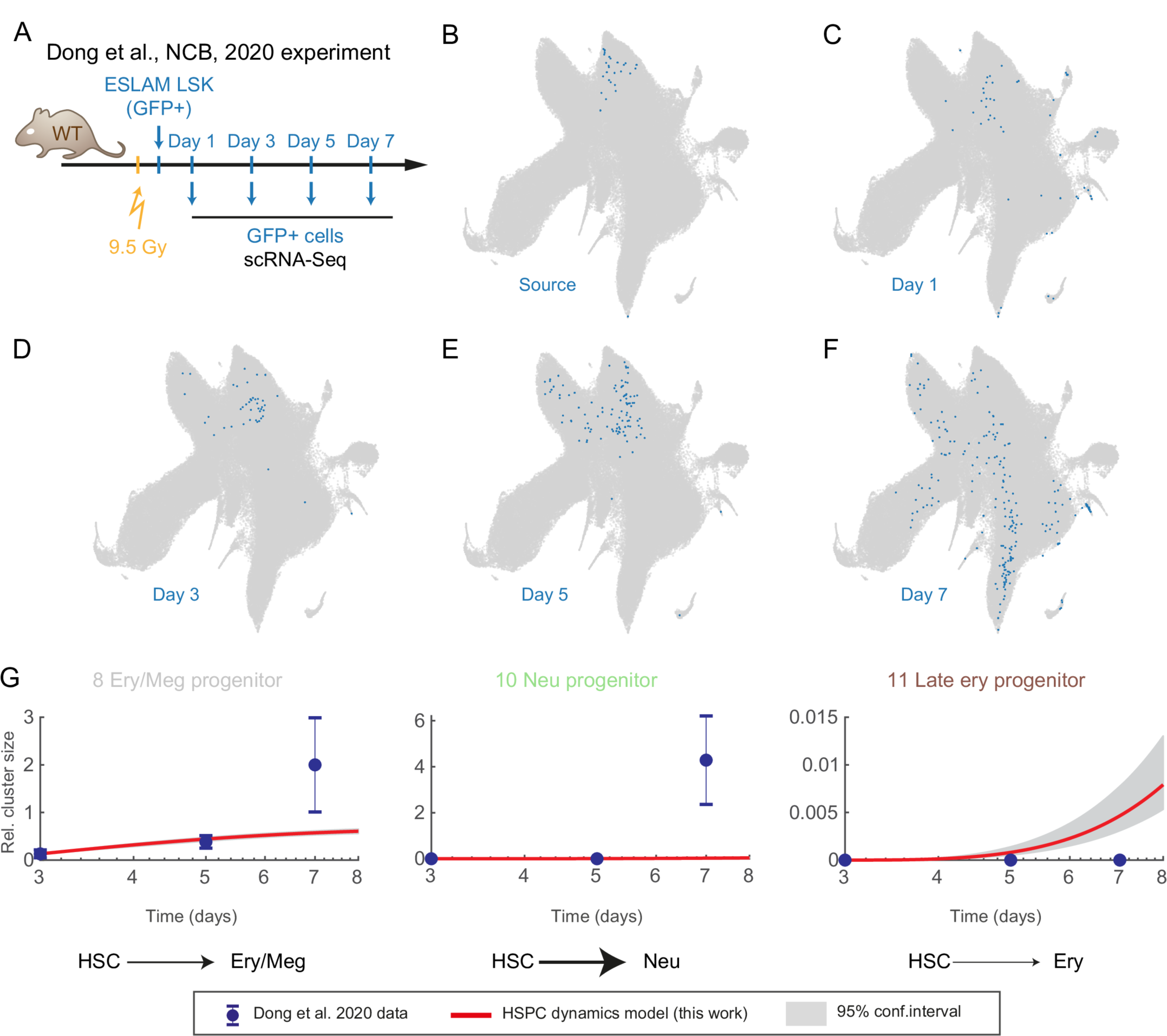
Growth and differentiation rates of HSPCs adapt to cellular stress conditions. (A) Diagram of the experiment performed by (Dong et al., 2020) study. (B-F) UMAP projections of the HSPC landscape (grey) with embedded cells from (Dong et al., 2020) in blue. (G) Relative cluster size, data (points) and discrete model prediction (red line with 95% confidence interval) based on day 3 data from (Dong et al., 2020). Error bars indicate propagated standard error of the mean.

## Discussion

Quantitative models describing cell differentiation (e.g. Waddington landscape) have been conceptualized decades ago (Waddington, 1957). Yet, we have barely progressed beyond static and qualitative abstractions of hematopoiesis. Here, we report a major effort which has allowed us to add real time to single-cell transcriptomics data. Analogously to the moving images in a Kinetoscope, our approach uses snapshots of differentiation to reconstruct the cellular flow between single-cell states within the bone marrow. Internally, our model describes cell behavior as a function of self-renewal and differentiation rate parameters, which in turn translate into the shape of a Waddington-like landscape (Figure 6). The discrete model approximates the landscape with a set of pre-defined platforms connected with slides, whereas the continuous model follows the shape for all observed states (here: single cells). Differentiation rate is analogous to the slope of the valley connecting two states, with steeper slopes indicating faster transition. In turn, stable states have little or no downward slope and combined with proliferation constitute areas of high self-renewal - these can be imagined as flat areas in the landscape (Figure 1G).

**Figure 6.**
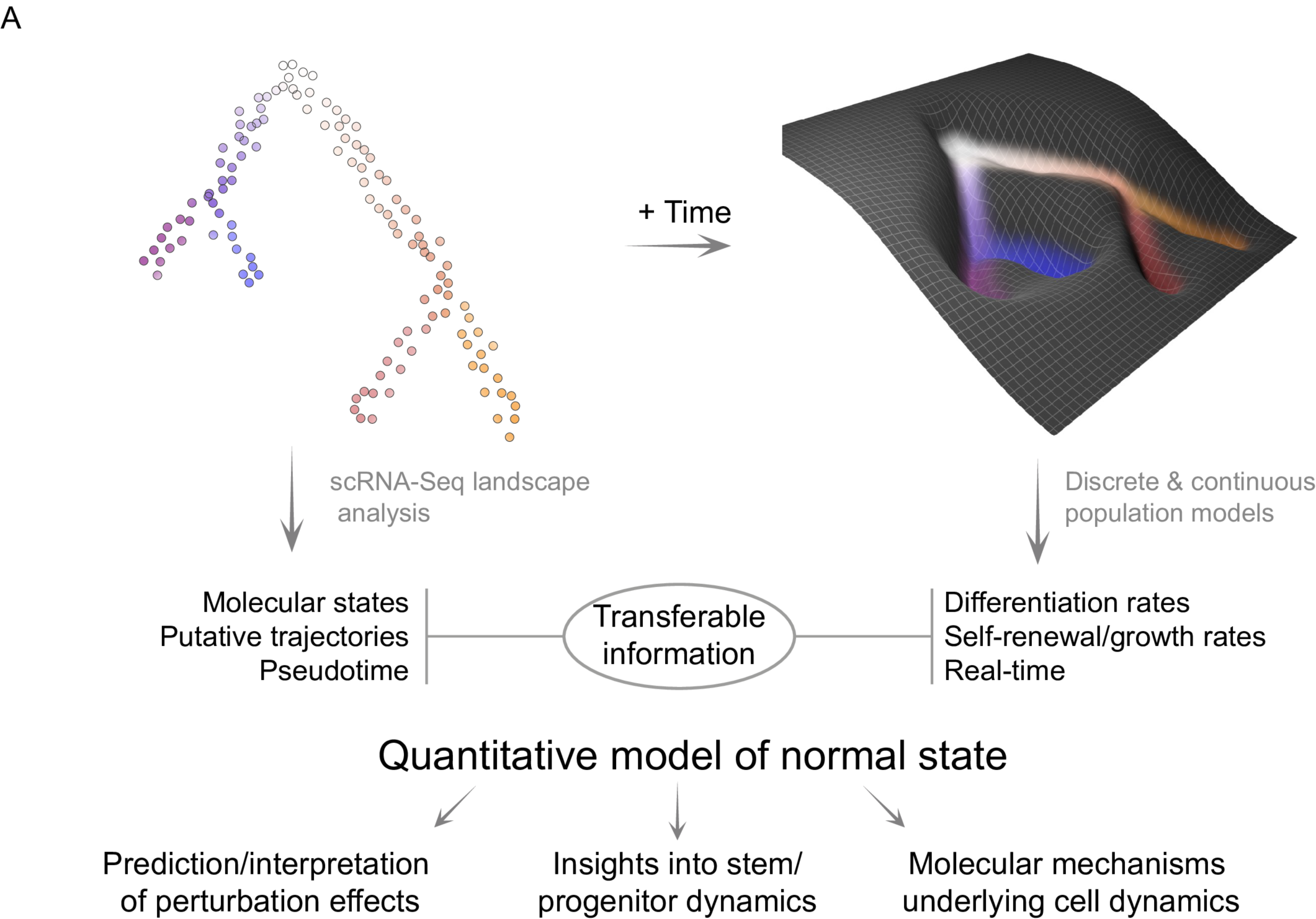
The quantitative model of HSPC dynamics in the mouse bone marrow. Diagram highlighting the transferable information and the model utility.

The cellular context is crucial for understanding the differentiation rates and cell fate. CMPs have been proposed as an intermediate progenitor population with combined erythroid, megakaryocytic, neutrophilic and monocytic potential (Akashi et al., 2000). However, later studies reported that the majority of CMPs are transcriptionally and epigenetically primed towards specific lineages (Paul et al., 2015), accompanied by lineage bias and mostly unipotent output (Perié et al., 2015) in transplantation cell fate assay. Importantly, transplantation, as we show in this work, is associated with greatly increased differentiation rates, most likely due to high proliferative demand, which if induced by 5-FU treatment also causes accelerated differentiation (Busch et al., 2015). In fact, cell fate assays performed in media containing a range of differentiation-supporting cytokines, rarely show combined megakaryocyte, erythroid, granulocyte and monocyte output, as demonstrated by (Akashi et al., 2000) and (Weinreb et al., 2020). Instead, if cells are given an opportunity to expand (for approx. 3 divisions) under cytokine-restricted conditions (SCF, IL-11, TPO only) >50% CMPs are capable of generating multipotent output in secondary assays (Akashi et al., 2000). The same argument applies to the LMPP population, which was originally reported as consisting mostly of multi-potent cells based on analogous two-phase culture assays (Adolfsson et al., 2005) and later described to produce mostly unilineage output in transplantation assays (Naik et al., 2013). Our model suggests that the clusters 8, 4, 5 (largely overlapping with CMPs) transition slowly among each other and particularly in case of cluster 8 and 4 both directions are permitted, indicating that these states may exist in balance. Thus, strong differentiation conditions (transplantation or differentiating media) are likely to simply not provide enough time for cells to ’explore’ the multipotent progenitor space. Moreover, if a progenitor cell does not divide before being channeled down a particular lineage, alternative fates can never be realized. Further work will be required to better resolve the HSC sub-populations (in cluster 0). We consider the tentative sub-structure provided here as a useful first step in this endeavor, as it fits both our data and experimental evidence of HSC behavior in ageing mice (Barile et al., 2020; Busch et al., 2015; Yamamoto et al., 2018; Zhang et al., 2020).

We fully leverage the scRNA-Seq approach to vastly extend our model’s applicability. To ensure accessibility and interpretability for wide audience we integrated published annotation from many sources. This places our unified landscape (and its sub-populations) in the biological context of previous immunophenotyping and lineage tracing experiments. Moreover, static cell properties (cluster, pseudotime) and model parameters (differentiation rates, self-renewal) are transferable. As demonstrated in Figure 5, new time-course scRNA-Seq data can be readily incorporated into our landscape, while the model highlight changes in cell differentiation rates in context of cell transplantation. Even with snapshot data our model can be used to simulate what changes in cell dynamic parameters could underlie an observed difference in cell abundance between conditions.

Differentiation and growth involve coordinated up- and down-regulation of thousands of genes, where it remains unknown for the vast majority of those genes whether and if so, how, they play a role in controlling cell behavior. To access the relevant molecular states with high precision, we introduce the first continuous model of native hematopoiesis which includes per-cell growth and differentiation rates, thus providing a direct comparison between cellular behavior and underlying gene expression. We observed complex, sequential gene expression pattern, some of which overlap with increasing differentiation rates, suggesting irreversible molecular changes. Specifically, we show that neutrophil differentiation is coupled with expression of multiple lineage determinants (*Irf8*, *Flt3*, *Pou2f2*, *Gfi1*) followed by a single programme taking over and a further increase in differentiation.

Our work shifts the paradigm from qualitative models with limited predictive capabilities to integrative and quantitative models. The latter are highly transferable and thus key to providing insight into human hematopoiesis, where experimental options are limited. Recently demonstrated scRNA-Seq information transfer across species (Lotfollahi et al., 2019, 2022; Welch et al., 2019) will potentially enable mapping HSPC dynamics onto human counterparts. Self-renewal and differentiation capacities are particularly relevant to leukemia research, which, in most cases, struggles with identifying cell types of cancer origin. As we show here and supported by previous studies (Busch et al., 2015; Takahashi et al., 2021), progenitors can also operate close to self-renewal and a small proliferative advantage may be sufficient to immortalize them. Finally, population dynamic models are universally applicable across biological fields, as adult tissues are commonly replenished from their own stem cell pools (Goodell et al., 2015). To instruct such future endeavours, we demonstrated how to build a model connecting high-resolution molecular information with tissue-scale cell behavior.

## Methods

### Data and code availability

- All sequencing data has been deposited on GEO under accession number: GSE207412.
- Cellproject package is available through Github (https://github.com/Iwo-K/cellproject).

### Hoxb5^CreERT2^ mouse line

The *Hoxb5^CreERT2^* allele was generated using CRISPR-Cas9 gene editing technology employing fertilized 1-cell zygotes on the B6CBAF1/Crl genetic background. We injected a single 15 ng/ul sgRNA (tcctccggatgggctca) (Chen et al., 2016) together with 25 ng/ul CAS9 mRNA and 17.5 ng/ul single strand donor DNA encoding the P2A-CRE-ERT2 protein flanked by 70 nucleotides of homology arms (Supplementary Table 2). The F0 offspring was screened by PCR and Sanger sequencing. The *Hoxb5^CreERT2^* line was established from one founder animal and back-crossed several times to the C57BL/6N genetic background. Mice were genotyped by PCR using primers detailed in Supplementary Table 2.

### Induction of reporter gene expression by tamoxifen

Tamoxifen (1g; Sigma T5648) was dissolved in 10 mL absolute ethanol and 90 mL corn oil (Sigma C8267) at 37°C. Aliquots of tamoxifen (10 mg/mL) were stored at -20 °C. Mice were injected intraperitoneally (i.p.) with tamoxifen at 100 mg/kg body weight for 7 days. Hoxb5-Tom mice were injected with equivalent volume of corn oil to confirm no background or tamoxifen-induced changes in the reporter strain alone. Non-injected Hoxb5-Tom mice were also analysed to determine whether any labelling was present in the absence of induction.

### Flow cytometry

At end point analyses, the fraction of Tom^+^ cells was determined in various hematopoietic compartments of BM, PB, spleens, thymi and lymph nodes. Cells from those tissues were prepared and analyzed as described previously (Lawson et al., 2021; Mapperley et al., 2021).

For HSC and progenitor cell analyses, unfractionated BM cells were incubated with Fc block (BD Pharmigen 553141), followed by biotin-conjugated anti-lineage marker antibodies (CD4 (Biolegend 553649), CD5 (Biolegend 553019), CD8a (Biolegend 553029), CD11b (Biolegend 553309), B220 (Biolegend 553086), Gr1 (Biolegend 553125) and Ter119 (Biolegend 553672)), cKit-BV711 (Biolegend 105835), Sca1-APC/Cy7 (Biolegend 108126), CD48-APC (Biolegend 103411) and CD150-PE/Cy7 (Biolegend 115914) antibodies. Biotin-conjugated antibodies were then stained with Pacific blue-conjugated streptavidin (Biolegend 405225). DAPI (BD Pharmigen 564907) was used for dead cell exclusion.

For analyses of differentiated cells in the BM, cell suspensions were stained with B220-APC (Biolegend103212) and CD19-APC/Cy7 (Biolegend 115529) antibodies for B cells, CD11b-PB (Biolegend 101224) and Gr1-PE/Cy7 (Biolegend 108416) for myeloid cells and Ter119-FITC (Biolegend 116206) for erythroid cells.

For analyses of differentiated cells in PB, spleen and lymph node, myeloid cells were stained as above for BM cells, T cells with CD8a-APC (Biolegend 100712) and CD4-APC (Biolegend 100411) antibodies, and CD19-APC/Cy7 (Biolegend 1155290) antibodies were used to detect B cells.

Cell suspensions from thymus were incubated with the biotin-conjugated anti-lineage marker antibodies described above together with CD4-APC (Biolegend 100411), CD8b-APC/Cy7 (Biolegend 126620), CD25-PB (Biolegend 102022) and CD44-PE/Cy7 (Biolegend 103030) antibodies. Biotin-conjugated antibodies were then stained with PerCP-conjugated streptavidin (Biolegend 405213). Flow cytometry analyses were performed using LSRFortessa (BD).

### Cell isolation for the scRNA-Seq experiments

All steps in this section (unless otherwise indicated) were performed on ice, and centrifugation steps performed at 300g, 4°C for 5 min. 8-12 weeks old mice carrying the Hoxb5-Cre and the Rosa26-LoxP-STOP-LoxP-tdTomato constructs were treated with 7 daily injections of tamoxifen (as described above) and sacrificed at indicated time-points. Bone marrow cells were extracted from ilia, tibiae and femora by grinding with mortar and pestle in PBS supplemented with 2% Fetal Bovine Serum (cell buffer). The suspension was filtered through a 50µm filter, centrifuged and resuspended in 3 ml of cell buffer. Red blood cells were removed using the ammonium chloride solution: 5 ml of 0.8% Ammonium Chloride (StemCell Tech. 07800) was added to the suspension and incubated for 10 min with intermittent mixing. Afterwards cells were diluted with 7 ml of cell buffer, centrifuged and resuspended in 1 ml of cell buffer. Subsequently, lineage depletion was performed as follows: added 20 µl of the EasySep mouse hematopoietic progenitor cell isolation cocktail (Stem Cell Technologies 19856), incubated for 15 min, added 30 µl magnetic particles, incubated for 10 min, added 1.5 ml of cell buffer and placed tubes in a magnet, incubated for 3 min at room temperature and eluted cells twice (with additional 2.5 ml of cell buffer). Afterwards, cells were centrifuged, resuspended in 200 µl of cell buffer and stained with the antibody panel as follows: antibody mix was added, cells were incubated for 30 min, washed with 2 ml of cell buffer, centrifuged, resuspended in 200 µl cell buffer. For the secondary staining Streptavidin-BV510 was added, cells were washed with 2 ml of cell buffer, centrifuged, and resuspended in 1000 µl of cell buffer supplemented with 7AAD. Afterwards cells were sorted with BD influx sorter into either 96 well plates containing 2.3 µl lysis buffer (for the Smart-Seq2 protocol) or 100 µl of PBS with 0.04% BSA in eppendorf tubes (’droplet buffer’) when used for the 10x Genomics scRNA-Seq protocol. The Smart-Seq2 plates were vortexed, centrifuged at 800g for 2 min and stored at -80°C.

Both Tom^+^ or Tom^-^ cells within the Lin^-^/(cKit OR Sca1)^+^ gate were sorted. (cKit OR Sca1)^+^ is a superset of the cKit+ gate used previously (Dahlin et al., 2018) which contains more lymphoid progenitors and pDCs.

Antibodies and fluorochromes used: Mouse hematopoietic progenitor cell isolation cocktail (Stem Cell Technologies 19856), CD48-APC (Thermo-Fischer 17-0481-82), c-Kit-APC/Cy7 (Biolegend 105826), Sca1-BV421 (Biolegend 108133) and CD150-PE/Cy7 (Biolegend 115914), Streptavidin-BV510 (Biolegend 405234), 7AAD (Thermo Fischer A1310)

### scRNA-seq data generation

#### Smart-Seq2

When cell numbers were limiting single cells were profiled with a modified version of the Smart- Seq2 protocol (Bagnoli et al., 2018; Picelli et al., 2014) rather than 10x Genomics kit. Single cells were sorted into 96-well plates with 2.3 µl lysis buffer containing 0.115 µl of SUPERase-In RNase Inhibitor at 20 U/µl concentration (ThermoFisher AM2694) and 0.23 µl of 10% Triton X-100 solution (Sigma 93443), plates were vortexed and stored at -80°C. After thawing 2 µl of the annealing solution (0.1 µl of ERCC RNA Spike-In solution (1:300,000 dilution) Thermo-Fisher 4456740), 0.02 µl of the oligo-dT primer (100 µM stock concentration) and 1 µl of dNTP (10 mM stock concentration)) was added. The plate was incubated at 72°C for 3 min, cooled down on ice and reverse transcription was performed by adding 5.7 µl of RT buffer (0.1 µl of Maxima H minus reverse transcriptase at 200 U/µl concentration (ThermoFischer EP0752), 0.25 µl of SUPERase- In RNAse Inhibitor at 20 U/µl concentration, 2 µl of the Maxima enzyme buffer, 0.2 µl of TSO oligo at 100 µM concentration, 1.875 µl of PEG 8000 solution (Sigma P1458) at 40% v/v concentration and 1.275 µl water) and incubation at 42°C for 90 min followed by incubation at 70°C for 15 min. Immediately after, cDNA was amplified by PCR by adding 1 µl of the Terra PCR Direct Polymerase (1.25 U/µl, Takara 639270), 25 µl of the Terra PCR Direct buffer and 1 µl of the ISPCR primer (10 µM stock concentration) to a total volume of 50 µl using the following PCR conditions: 98°C for 3 min, 98°C for 15 s, 65°C for 30 s, 68°C for 4 min (21 cycles), 72°C for 10 min. The amplified cDNA was purified using AMPure XP beads (Beckman A63882), quantified using the PicoGreen assay (ThermoFischer P7589) and used for Nextera library preparation. The libraries were generated using either a standard protocol (batch 7d) or modified protocol (batches 3d7d, 2w4w and 3dr2), see the corresponding metadata) described below. No obvious batch effects were observed among cells analyzed with either of the protocols.

The standard Nextera protocol: cDNA was diluted to approximately 50-150 pg/µl and 1.25 µl of the solution was used, 2.5 µl of Tagment DNA buffer 1.25 µl of Amplicon Tagment Mix (Nextera XT kit, Illumina FC-131-1096) were added, samples were incubated at 55°C for 10 min, and the reaction was stopped by addition of 1.25 µl of NT buffer. Tagmentation products were amplified by PCR by adding 1.25 µl of each N and S primers (Illumina Nextera XT 96-index kit FC-131- 1002) and 3.75 µl of NPM solution and using the following thermocycler settings: 72°C 3 min, 95°C 30 s, 12 cycles of 95°C 30s, 55°C 30s, 72°C 60s and a final extension at 72°C for 5 min.

The modified Nextera protocol follows the same principle as the standard Nextera protocol and includes the following steps: cDNA was diluted to approximately 50-150 pg/µl and 1.03 µl of the solution was used, 1.63 µl of Tagment DNA buffer and 0.6 µl Amplicon Tagment Mix was added, samples were incubated at 55°C for 10 min, the reaction was stopped by adding 0.82 µl of NT buffer. Tagmentation products were amplified by adding 1.23 µl of each N and S primers (as above but diluted 5 times), 2.3 µl of Phusion HF buffer (ThermoFischer F530L), 0.1 µl of dNTP (25 mM stock concentration), 0.07 µl of Phusion polymerase and 2.5 µl of water and using the following thermocycler settings: 72°C 3 min, 98°C 3 min s, 12 cycles of 98°C 10s, 55°C 30s, 72°C 30s and a final extension at 72°C for 5 min.

Libraries were sequenced using the Illumina Hiseq4000 or NovaSeq instruments, obtaining approximately 100-200 mln reads per 96 cells.

### 10X genomics

For the 10x Genomics scRNA-Seq protocol up to 20,000 cells were pooled in pairs corresponding to male and female animals, centrifuged and resuspended in a volume of droplet buffer optimal for recovery of up to 10,000 cells and immediately processed with the 10x Genomics Single Cell 3’ v3 protocol following the manufacturer’s instructions.

Libraries were sequenced using the Illumina NovaSeq instrument, obtaining at least 20,000 reads per cells.

### scRNA-Seq data analysis

Smart-Seq2 sequencing reads were aligned to the mouse genome (mm10) using the STAR aligner (version 2.7.3a) with default parameters. Reads mapping to exons were counted with featureCounts (version 2.0.0) using the ENSEMBL v93 annotation. Each sample was subjected to a quality control, samples with: <100,000 reads, <23% of reads mapped to exons, >8.5% of reads mapped to ERCC transcripts, >10% mitochondrial reads or <2000 genes detected above 10 counts per million were discarded. 1288 out of 1533 samples passed quality control. Data were normalized 10,000 total counts and ln(n+1) transformed.

10x genomics reads were pre-processed using cellranger (version 3.1.0, reference genome and annotation version 3.0.0) with default settings. Downstream analysis was performed mainly using the scanpy (Wolf et al., 2018) framework with additional packages where indicated. Low quality barcodes with less than 1000 genes were excluded from the analysis, doublet scores were estimated using the scrublet tool (using 30 principal components), potential doublets were removed. Male and female cells were distinguished based on the expression of the Xist gene and Y chromosome genes. Cells with detectable Xist expression and undetectable Y chromosome gene expression were classified as female and vice versa, ambiguous cells or potential doublets were excluded. Data were normalised to 10,000 total counts and ln(n+1) transformed.

To determine highly variable genes, scanpy’s highly_variable_genes function was used to select top 5000 genes within the 10x genomics data. From the list of highly variable genes, genes associated with cell cycle, Y-chromosome genes and the Xist were excluded. Genes associated with cell cycle were a union of cell-cycle genes from (Dahlin et al., 2018) and genes with at least 0.1 Pearson correlation with the following gene set: Ube2c, Hmgb2, Hmgn2, Tuba1b, Ccnb1, Tubb5, Top2a, Tubb4b, following the method from (Weinreb et al., 2020). Putative cell cycle phase was assigned using scanpy’s ’score genes cell cycle’ function to assign putative cell cycle phase to both 10x and Smart-Seq2 cells. Following that, 10x and Smart-Seq2 data were combined and subjected to Seurat CCA batch correction (Stuart et al., 2019). Among a variety of batch correction tools (Harmony (Korsunsky et al., 2019), Scanorama (Hie et al., 2019), BBKNN (Polański et al., 2020), fastMNN (Haghverdi et al., 2018), MNNcorrect) only Seurat CCA generated seamless integration best matching the cell frequencies based on flow cytometry analysis. After applying batch correction, we observed no obvious segregation of Smart-Seq2 and 10x scRNA-Seq profiles (Figure S2E). Corrected log-normalized counts were scaled and used to compute 50 principal components, find nearest neighbors and calculate a UMAP projection (McInnes et al., 2020). A minor batch effect between 10x samples was corrected using Harmony batch correction tool (Korsunsky et al., 2019). The corrected principal components were used to calculate 12 neighbors followed by cell clustering using the leiden algorithm (Traag et al., 2019) and calculation of the UMAP projection. Clusters were manually annotated based on the marker gene expression as described in Supplementary table S1. To reduce the complexity for the discrete model clusters with the following criteria were excluded from the further analysis: clusters that appeared disjointed from the main landscape body, represented low-quality/dying cells or with unclear origins based on the UMAP projection and PAGA analysis. This included: T cells, innate lymphoid cells (ILCs), cells with high mitochondrial gene counts, mature B cells, interferon-activated cells, cells with high complement expression and small clusters with unclear annotation, likely to represent doublet cells. Unfiltered landscape is displayed in Figure S2F,G.

To visualize the relative proportions of cells per cluster over time (Figure S3B), we averaged fractions of Tom^+^ cells in each cluster for each time-point and divided by the respective values for matching Tom^-^ cells.

### Embedding external datasets into the integrated HSPC landscape

For each external datasets the log-normalised counts for cells passing quality control were used as in the original work. Annotation was either obtained from the respective GEO repositories, literature or kindly provided by the authors.

Each dataset was integrated with the HSPC landscape (below denoted as reference) using the indicated batch correction tools and the Cellproject package as follows. Log-normalized counts for (Nestorowa et al., 2016) were concatenated with the reference and batch effect was removed using Seurat CCA method (Stuart et al., 2019) only highly-variable genes selected in the reference landscape were used. The corrected values were scaled and used to compute PCA (50 components) in the reference dataset. The correct values of (Nestorowa et al., 2016) dataset were fit into the reference PCA space, in which 15 nearest neighbors were identified between the datasets. These nearest neighbors were used for two purposes: (1) transfer the cluster identity to the new data (based on the most frequent label) and (2) to predict coordinates in the original reference PCA space (used as a basis for UMAP projection) using nearest-neighbor regression. Finally, the new PCA coordinates were used to embed the new data into UMAP space. As immunophenotypic populations we used the ’narrow’ classification provided in the original study.

(Bowling et al., 2020) data was concatenated with the reference and a common PCA space was calculated, which was subsequently corrected with the Harmony batch correction tool. Within the corrected space 8 nearest neighbors were identified across the datasets, followed by label transfer and UMAP embedding as described above.

(Weinreb et al., 2020) data was integrated analogously to the (Nestorowa et al., 2016) data. Only ’state-fate’ clones were used, ie. cells captured at an early time-point (day2) with measured fate outcomes at later time-points. Only fates with more than 7 cells were considered for the analysis.

### Trajectory inference and selection

To pinpoint the most immature stem cells the HSC score was calculated (default parameters) (Hamey and Göttgens, 2019) and denoised by averaging values over the nearest neighbors for each cell. As diffusion pseudotime the cell with the highest smoothed HSC score was selected, diffusion map was calculated and served as the basis for trajectory inference and continuous populations models described below.

To infer putative trajectories Tom^+^ cells were used (matching the Pseudodynamics analysis below) for calculating cell transition probabilities using the Pseudotime Kernel method (based on the Palantir tool (Setty et al., 2019)) from the CellRank package (Lange et al., 2022). To define the end states clusters 6, 7, 10, 11, 13, 14, 15, 16, 17, 18, 19 were selected and within them 50 cells with the highest pseudotime values. These states are largely consistent with an unsupervised method of macrostate selection Generalized Perron Cluster Analysis with Schur decomposition (Lange et al., 2022). To assign cell fate probabilities Cellrank’s compute_absorption_probabilities function was used.

Cells belonging to trajectories for the continuous models were selected as follows. In case of megakaryocytic trajectory cells belonging to cluster 0, 7 and 8 and with the respective fate probability >0.3 were chosen. For the erythroid trajectory cells with respective fate probability <0.2 and falling within the pseudotime range 0.015 and 0.294 (to exclude variable small number at the end of the trajectory) were used. Neutrophil and monocyte share a long stretch of progenitors with high probabilities towards both lineages, thus a different approach was used, motivated the apparent locations of bipotent cells with neutrophil and monocyte/DC potential based on cell fate assays (Figure 2G) (Weinreb et al., 2020). Neutrophil progenitors (terminal state 10) were selected with fate probability >0.24 and Mono/DC probability <0.38 and excluding a small number of cells falling into clusters 12, 17 and 14. Conversely for the Mono/DC progenitors (terminal state 6) cells were selected with Mono/DC fate probability >0.18 and neutrophil probability <0.49 and a small number of cells falling into clusters 12, 17 and 14 was excluded.

### Discrete population model analysis

As input to the discrete models the estimated total number of Tom^+^ or Tom^-^ cells per cluster was used. The numbers were estimated based on the fraction of cells assigned to each cluster adjusted by the total number of cells analyzed by flow cytometry (samples were analysed in their entirety). One out of 5 mice analyzed at day 3 exhibited abnormally high labelling frequency, the sample was excluded to avoid introducing bias but we provide the corresponding data within the GEO submission files and source code for individual assessment.

To assess the kinetics of differentiation and growth of the different hematopoietic populations, we first considered a discrete compartments model, using the HSPC landscape clusters as compartments. To establish the topology of the differentiation process, PAGA connections and pseudotime ordering were considered. First of all, only transitions with PAGA connectivities higher than 0.05 were selected. No back differentiation (ie. against pseudotime ordering) was permitted into cluster 0 and from most differentiated clusters with clear expression of commitment genes: 1, 3, 6, 7, 9, 10, 11, 12, 13, 14, 15, 16, 17, 18, 19. Other transitions above-threshold were considered potentially bidirectional. Each compartment is assigned a growth rate and as many differentiation rates as the number of its progeny compartments. Assuming the following:

- the label is neutral and stably propagated
- the kinetics parameters of each cluster are constant over time and independent of the size of any cluster
- the labeled and unlabeled cells have identical kinetics,

Population dynamics can be modelled as an ODE system of coupled equations:

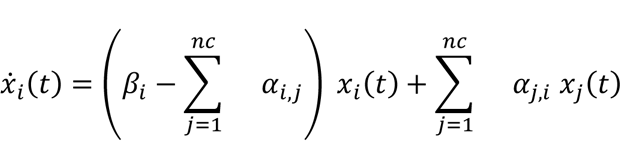

where *x*_*i*_(*t*) is the number of cells in population *i*, *α*_j,i_ is the differentiation rate from compartment *j* to *i*, and *β*_*i*_ the growth rate of population *i*. For the terminal and initial clusters the equations take form respectively:

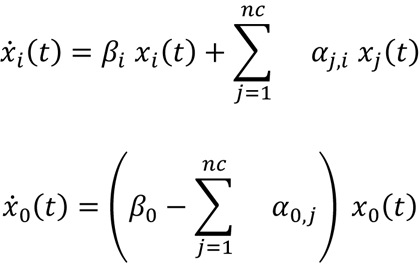

Please note that differentiation rates are set to zero if they have not passed the thresholding criteria as explained above. The number of clusters, nc, is equal to 22, one per each of the 20 Leiden clusters, plus 2 additional subpopulations within cluster 0, the most immature cluster. The reason for this choice lays in 2 observed characteristics in the data: cluster 0 ratio of labelled to unlabeled cells (labelling frequency) grows over time, and some downstream clusters’ labelling frequency overshoots the one in cluster 0. Based on (Barile et al., 2020) and (Takahashi et al., 2021), this implies that the progenitor cluster must be heterogeneous. Indeed, the most immature HSCs occupy only the tip of cluster 0 (Figure 2C). Particularly, we chose to add 2 more sub-compartments to allow for differentiation bias in the HSCs. The differentiation rates were allowed to vary between 0 and 4 per day, with the exception of cluster 0a’s rates, which were bounded to vary between 0 and 0.02 per day, based on previous knowledge of HSCs low activity (Barile et al., 2020; Oguro et al., 2013). The growth rates were bounded between -4 and 4 per day, to allow for death rate (negative values) or additional differentiation towards more mature cell states outside the presented HSPC landscape, or cell migration. The growth rate in cluster 0a was fixed in such a way to balance the differentiation rates, given the a priori knowledge that pure functional haematopoietic stem cells show only limited growth over time (Zhang et al., 2020). Furthermore, we observed that the total number of cells in cluster 0 plateaus as the mice age, similarly to what was previously observed for the HSC and MPP populations (Barile et al., 2020). We accounted for this upon modelling cluster 0 overall number of cells with a logistic function, and thus added a logistic parameter *ρ* and a carrying capacity *K*. Both parameters are positive and unconstrained. Specifically, we implemented the following equations for cluster 0a:

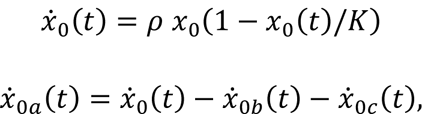

while the time evolution of clusters 0b and 0c is analogous to that of all other clusters. Since we calibrated the ODE system to both the labelled and unlabeled cells time courses, we also included as parameters 22*2 initial conditions, all positive and unbounded, except for the number of cells in cluster 0a, set to range between 500 and 1500 based on previous HSC number estimates (Kent et al., 2009) and factoring in cell isolation efficiency. The model allows the initial number of labelled cells to be greater than zero, thus accounting for any unspecific labelling.

We calibrated our model to 4 types of observables:

- The number of labeled cells in each cluster over time and relative to cluster 0 as computed via scRNA-Seq analysis
- The number of unlabeled cells in each cluster over time and relative to cluster 0 as computed via scRNA-Seq analysis
- The number of labeled cells in cluster 0 over time as computed via FACS sorting and scRNA-Seq analysis
- The number of unlabeled cells in cluster 0 over time as computed via FACS sorting and scRNA-Seq analysis

To estimate the parameters, we minimized a cost function of the squared sum of residuals. Each residual is weighted by the squared error, which was computed as pooled variance per time course. We computed the 95% confidence bounds on the parameters’ best fit with the profile likelihood method as in (Barile et al., 2020; Raue et al., 2009). To compute error bounds on the model, we ran ≈4000 bootstrap simulations, where data is resampled with replacement per time-point, and the cost function is re-minimized on the new dataset. For each simulation, a new parameter vector is found, and a model curve generated. 95% bootstrap confidence bounds are then determined cutting upper and lower 0.025 quantiles per time-point. To simulate the ablation of any population, the initial condition of the unlabeled cells for the corresponding compartment can be set to 0. To ablate the HSCs, we simultaneously set to 0 the initial condition of all 3 subclusters.

To compute the journey times, we generated the model in the time interval 1-300 days with 1 day steps, assuming that cells are initially only in cluster 0 and with the unlabeled cells initial condition. We then computed the smallest time for which the number of cells in a population reaches one and dubbed that journey time.

### Continuous population model analysis

In order to compute pseudotime-dependent kinetic rates, we relied on the pseudodynamics framework (Fischer et al., 2019). Briefly, the compartment model explained in the previous section has a one to one correspondence to the continuous model if the compartment index is treated as a continuous variable, namely the diffusion pseudotime coordinate *s*, the number of cells is replaced by the cell density over pseudotime and real time *u*(*s,t*), and the differentiation and net proliferation rates are replaced by the drift *v*(*s*) and the growth rate *g*(*s*), respectively. Given these substitutions, the ODE system becomes a PDE system. In addition, the Pseudodynamics framework also introduced an extra parameter *D*(*s*) that allows for diffusion of the cells on the pseudotime axis to account for stochasticity in the differentiation process. The 3 kinetics parameters, drift, growth rate and diffusion, are modelled as natural cubic splines with 9 nodes. The nodes boundaries were kept as in the original publication: between 0 and 1 per day for drift and diffusion, and between -5 and 6 per day for the growth rate. To simplify the computation, we estimated such rates independently for 4 different trajectories, which avoids introducing parameters that describe the branching process. The trajectories were chosen based on the affinity to each terminal state as estimated by CellRank (see section ’Trajectory inference and selection’). For each trajectory, the PDE reads:

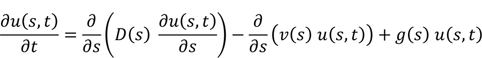

For the boundaries, we assumed no-flux Robin conditions, as in the original publication. To solve the PDE, we used the non-branching pseudodynamics model as compiled in MATLAB 2017b, with only one difference: we did not enforce differentiation to be 0 at the end of the trajectory which, together with the growth rates taking also negative values, accounts for the fact that the populations in our landscape are all transient and that fully mature cells are not captured by our gating strategy. The model was calibrated to the time-dependent density and total number of labelled cells only. The error was computed as variance among replicates. For each trajectory, at least 240 simulations were launched, with regularization parameters 0, 1, or 10 to penalize big differences in the splines’ nodes. The solution was chosen based on the highest log-likelihood, and the regularization parameter as the highest that visually fits the data well.

### Differential expression analysis

For the DE analysis cells were selected to match the continuous model trajectories. The shapes of differentiation and net proliferation rates were inspected for potential regions of interests and respective ranges of pseudotime values were chosen. Prior to the analysis genes with low expression were filtered out, only genes detected in more than 2.5% cells and with overall mean expression above 0.05 (data normalized with logNormCounts from the scuttle package) were included. To select genes with dynamic expression in the chosen intervals the fitGAM function followed by startVsEndTest from the TradeSeq package were used. Genes were considered significant if they showed at least FDR of 0.1 and a log2(Fold change) of at least 1. Predicted and smoothed gene expression was used using the predictSmooth function from the same package. In heatmaps genes were clustered with hierarchical clustering using the hclust R function with default settings. Transcription factors were selected based on the gene list established in (Ravasi et al., 2010), TF groups were established by cutting the tree at the level of 4. Gene enrichment was performed using GSEAPY interface to the enrichr tool (Kuleshov et al., 2016).

### Transplantation data analysis

(Dong et al., 2020) data was integrated into the HSPC landscape analogously to the (Nestorowa et al., 2016) data integration described in section ’Embedding external datasets into the integrated HSPC landscape’. Cells in each HSPC cluster were counted and used as an input into the discrete model prediction. Day 3 data was used as the initial condition and cell abundances per cluster were predicted from day 3 to day 7. The bootstrap confidence bounds were recomputed upon substituting the initial conditions. Given that the experimental data in relevant clusters vastly exceed the model prediction bounds, we concluded that the dynamics of perturbed haematopoiesis are different from normal conditions and suggest increased differentiation.

## Author contributions

### Part1 - Overall project design and Hoxb5-Tom model generation and characterization

Conceptualization K.R.K. and D.O.C.; Methodology J.C., F.S., N.B., P.N.M, K.R.K. and D.O.C; Software M.B.; Validation J.C., F.S., N.B., P.N.M., L.A., H.L., K.R.K. and D.O.C.; Formal Analysis I.K, J.C., M.B., F.S., N.B., K.R.K., D.O.C. and B.G.; Investigation J.C., F.S., N.B., P.N.M., L.A. and H.L.; Resources J.C., F.S., N.B., P.N.M., L.A., H.L., K.R.K. and D.O.C.; Data Curation J.C., F.S., N.B., L.A., H.L., K.R.K. and D.O.C.; Writing - Original Draft I.K., Writing - Review & Editing I.K., J.C., M.B., K.R.K., D.O.C. and B.G.; Visualisation I.K., J.C., M.B., F.S., N.B., K.R.K., D.O.C. and B.G.; Supervision H.L., K.R.K., D.O.C. and B.G.; Project Administration J.C., F.S., N.B., P.N.M., L.A., H.L., K.R.K., D.O.C. and B.G; Funding Acquisition K.R.K. and D.O.C.

### Part2 - scRNA-Seq analysis and HSPC dynamics modelling

Conceptualization I.K, M.B. and B.G.; Methodology I.K., M.B. and B.G.; Software I.K. and M.B.; Validation I.K., M.B. and B.G.; Formal Analysis I.K., M.B. and B.G.; Investigation I.K., J.C., N.B., M.L.R.H. and S.J.K; Resources I.K., J.C., M.B., K.R.K., D.O.C. and B.G.; Data Curation I.K., M.B and B.G.; Writing - Original Draft I.K. and M.B.; Writing - Review & Editing I.K, M.B., K.R.K., D.O.C. and B.G.; Visualisation I.K., M.B. and B.G.; Supervision I.K and B.G.; Project Administration I.K., J.C., M.B. and B.G; Funding Acquisition K.R.K. and B.G.

## Acknowledgements

The authors thank Reiner Schulte, Chiara Cossetti and Gabriela Grondys-Kotarba from the Cambridge Institute for Medical Research Flow Cytometry Core facility for their assistance with cell sorting. We would also like to thank Katarzyna Kania and others at the Cancer Research UK Cambridge Institute Genomics Core Facility for generating the 10x Genomics libraries and performing high-throughput sequencing. The authors are also grateful to all staff of the Biological Services Unit at Queen Mary University of London for their technical support. Work in the Kranc Laboratory is supported by Cancer Research UK (awards C29967/A14633 and C29967/A26787), The Barts Charity, Blood Cancer UK, and the Kay Kendall Leukaemia Fund. The O’Carroll laboratory is supported by the Wellcome Trust Investigator Award (106144), the Wellcome Centre for Cell Biology (203149) and a Wellcome multi-user equipment grant (108504). Work in the Göttgens laboratory is supported by Wellcome (206328/Z/17/Z and 203151/Z/16/Z), Blood Cancer UK (18002), Cancer Research UK (C1163/A21762) and UKRI Medical Research Council (MC_PC_17230). For the purpose of open access, the author has applied a CC BY public copyright licence to any Author Accepted Manuscript version arising from this submission.

---

**Figure S1.**
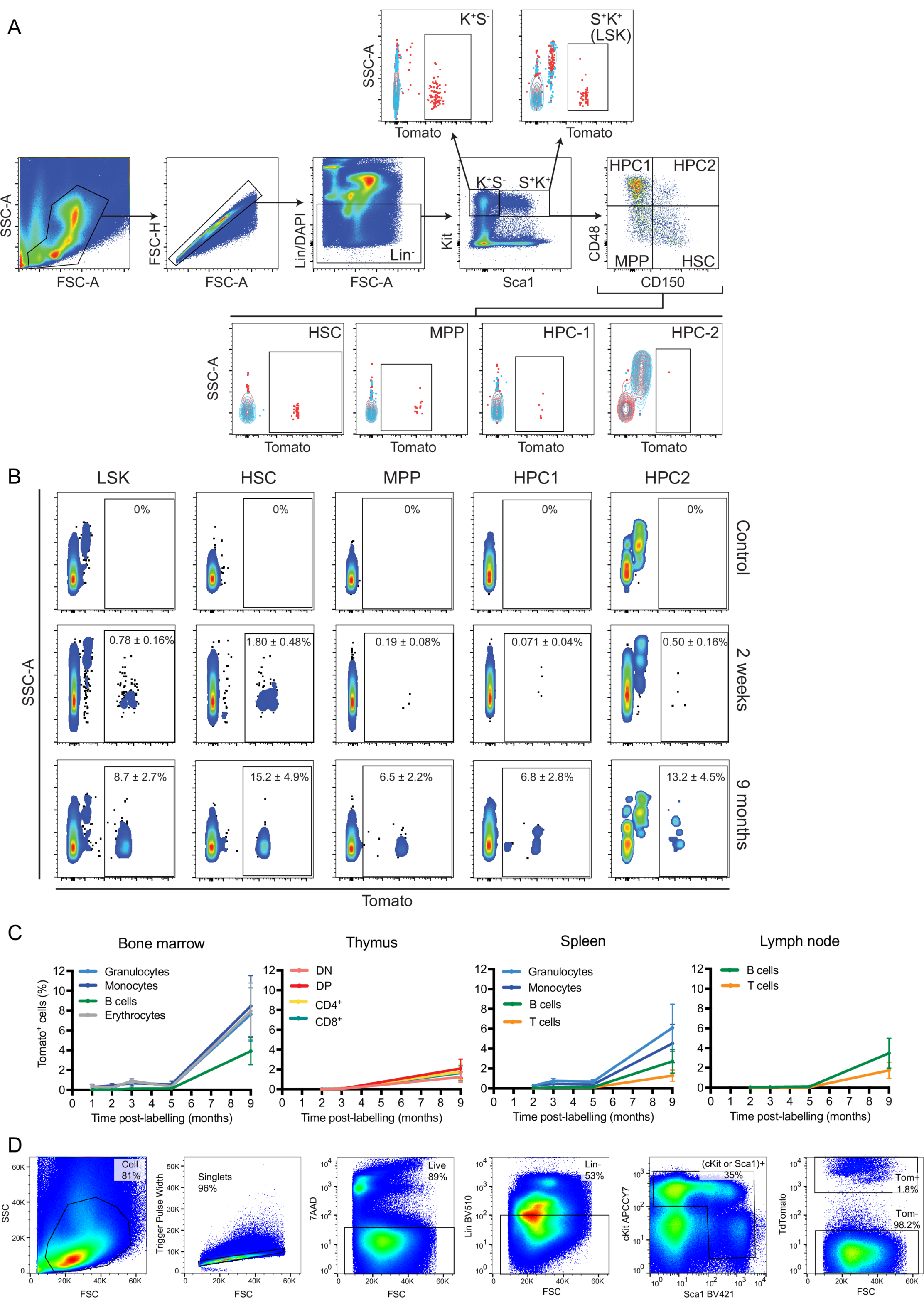
(A) Representative flow cytometry gates used for isolation of HSPC subpopulations and Tom^+^ cells from mouse bone marrow. Tom labelling (red) is shown in each population compared to control cells (blue). FACS plots correspond to mouse analysed 3 months after label induction. Relevant to Figure 1B,C. (B) Example flow cytometry plots showing Tom^+^ fractions in the bone marrow HSPC subpopulations from Hoxb5-Tom mice. Plots correspond to animals analyzed at 2 weeks and 9 months after label induction. Tom gate was set based on the signal from the control mice lacking the Tom label (top row). Relevant to Figure 1B, C. (C) Fractions of Tom ^+^ cells in the bone marrow, thymus, spleen and lymph nodes analyzed at the indicated time-points after label induction. Shown as mean with error bars denoting SEM of 4-32 animals. (D) Flow cytometry gating scheme for the Lin-, (Sca1^+^ OR cKit^+^) population (relevant to Figure 2 onward).

**Figure S2.**
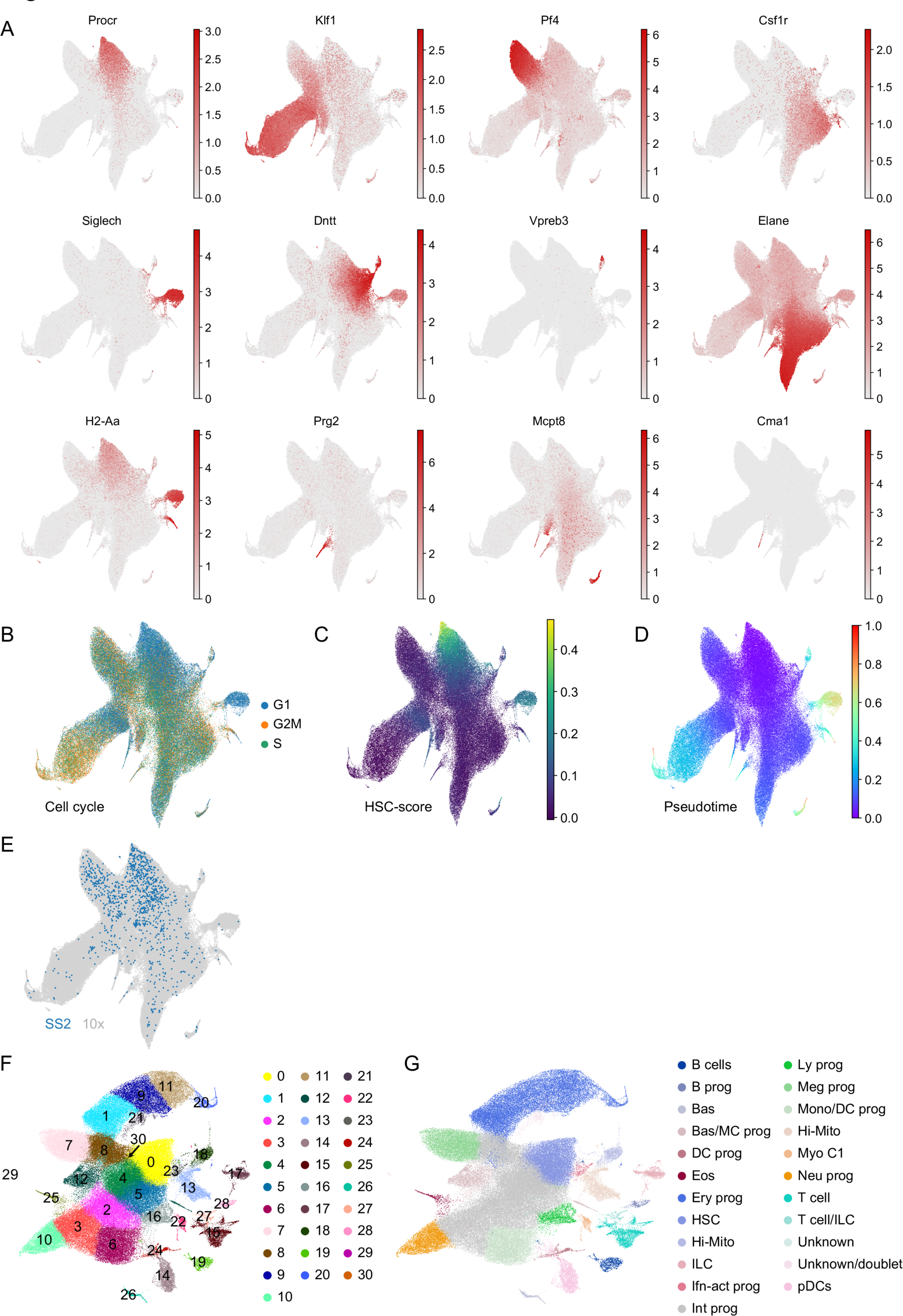
(A) UMAP projection of the integrated HSPC scRNA-Seq landscape (all Tom^+^ and Tom^-^ combined) with log-normalized expression for chosen marker genes in red. (B) Projection from A with inferred cell cycle phases. (C) Projection from A showing the HSC-score, metric correlated with the highest HSC repopulation potential. (D) Projection from A showing inferred diffusion pseudotime values for each cell. (E) Projection from A color-coded by scRNA-Seq technology used, SS2 - Smart-Seq2, 10x - 10x Genomics 3’ Kit. (F) UMAP projection of the integrated scRNA-Seq landscape (all Tom^+^ and Tom^-^ cells combined) prior to filtering out outlier/aberrant clusters with color-coded cluster information. After filtering cluster were renumbered in the consecutive order. (G) Projection from F with color-coded manual annotation. Abbreviations: B prog - B cell progenitor, Bas - basophils, Bas/MC prog - Basophil and Mast Cell progenitors, DC prog - dendritic cell progenitors, Eos - eosinophils, Ery prog - erythroid progenitors, HSC - hematopoietic stem cells, Hi-Mito - cluster characterized by high mitochondrial gene expression (potentially dying cells), ILC - innate lymphoid cells, Ifn-act prog - progenitors with strongly activated Interferon signature, Int prog - intermediate progenitors, Ly prog - lymphoid progenitors, Meg prog - megakaryocyte progenitors, Mono/DC prog - monocyte and dendritic cells progenitors, Myo C1 - myeloid cells with high expression of complement genes, Neu prog - neutrophil progenitors, pDC - plasmacytoid dendritic cells

**Figure S3.**
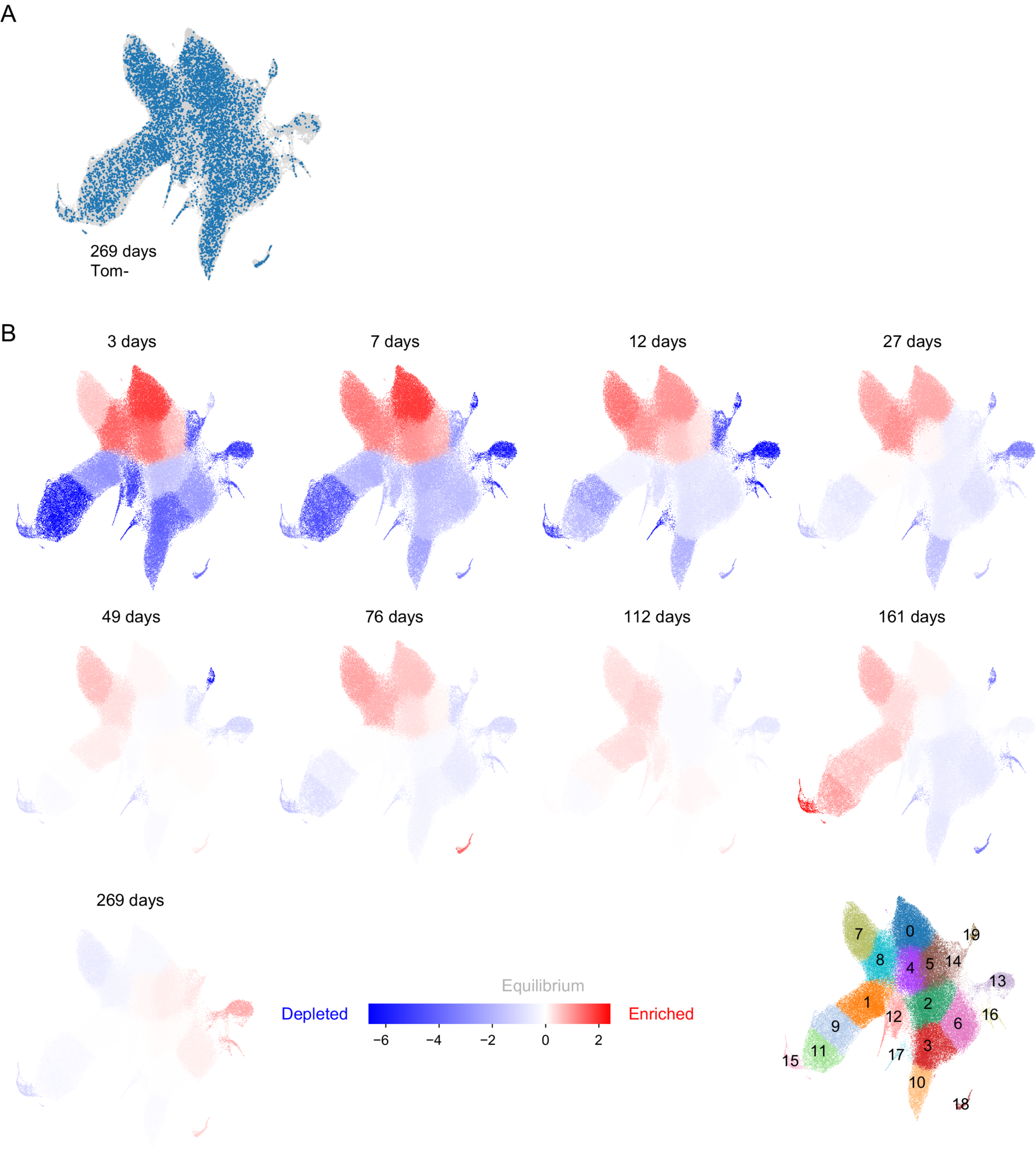
(A) UMAP projection of the integrated scRNA-Seq landscape (all Tom^+^ and Tom^-^ combined) in grey with Tom^-^ cells harvested at 269 days in blue. (B) Projection from A with each cluster color-coded according to its log2-transformed abundance ratio between Tom^+^ and Tom^-^cells. Relative abundance has been averaged across all samples for each time-point. Red indicates enrichment, white the expected value and blue depletion. For reference the cluster boundaries are visualized in the bottom right panel.

**Figure S4.**
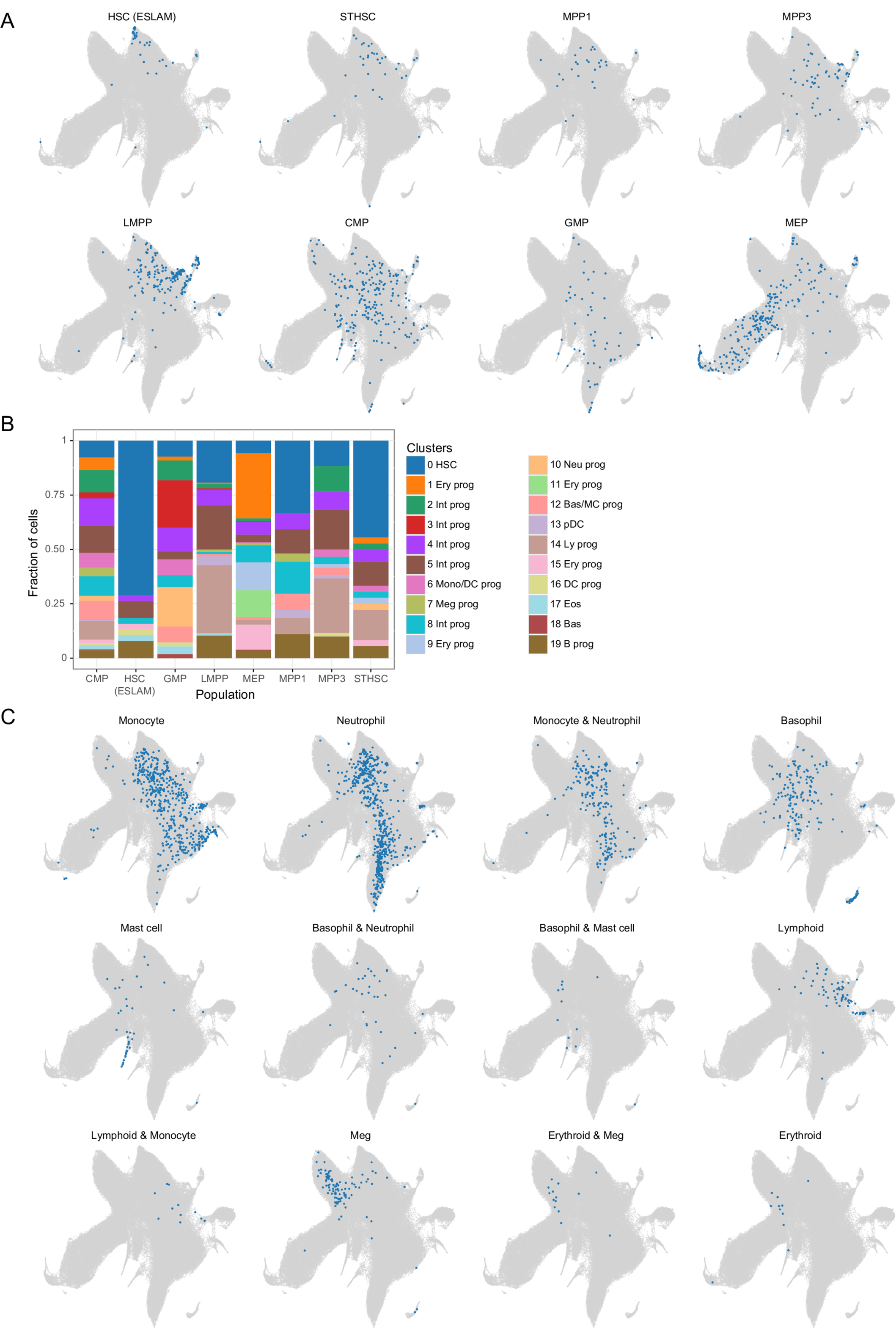
(A) UMAP projection of the integrated HSPC landscape (grey) with embedded immunophenotypic sub-populations (blue) from (Nestorowa et al., 2016) (B) Fraction of cells in each immunophenotypic population from A assigned to the HSPC landscape clusters. (C) UMAP projection of the integrated HSPC landscape (grey) with embedded cells from (Weinreb et al., 2020) split by their progeny fate.

**Figure S5.**
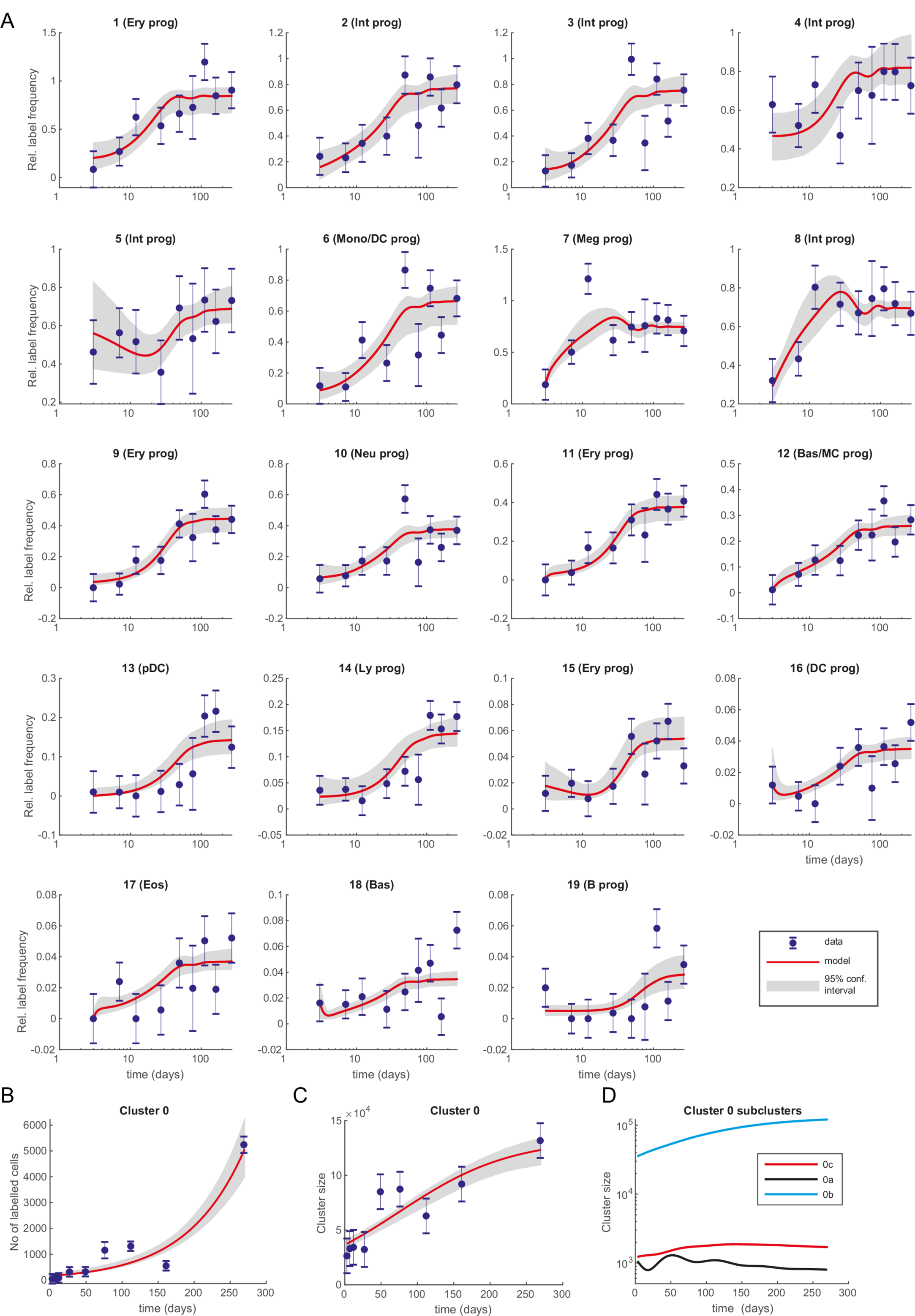
(A) Best discrete model fit (with 95% confidence intervals) for relative Tom^+^ label frequency by cluster (normalized to cluster 0). (B) Best discrete model fit (with 95% confidence intervals) for number of Tom^+^ cells in cluster 0. (C) Best discrete model fit (with 95% confidence intervals) for cluster 0 size. (D) Best discrete model fit for sub-cluster sizes within cluster 0. Error bars indicate pooled standard error of the mean.

**Figure S6.**
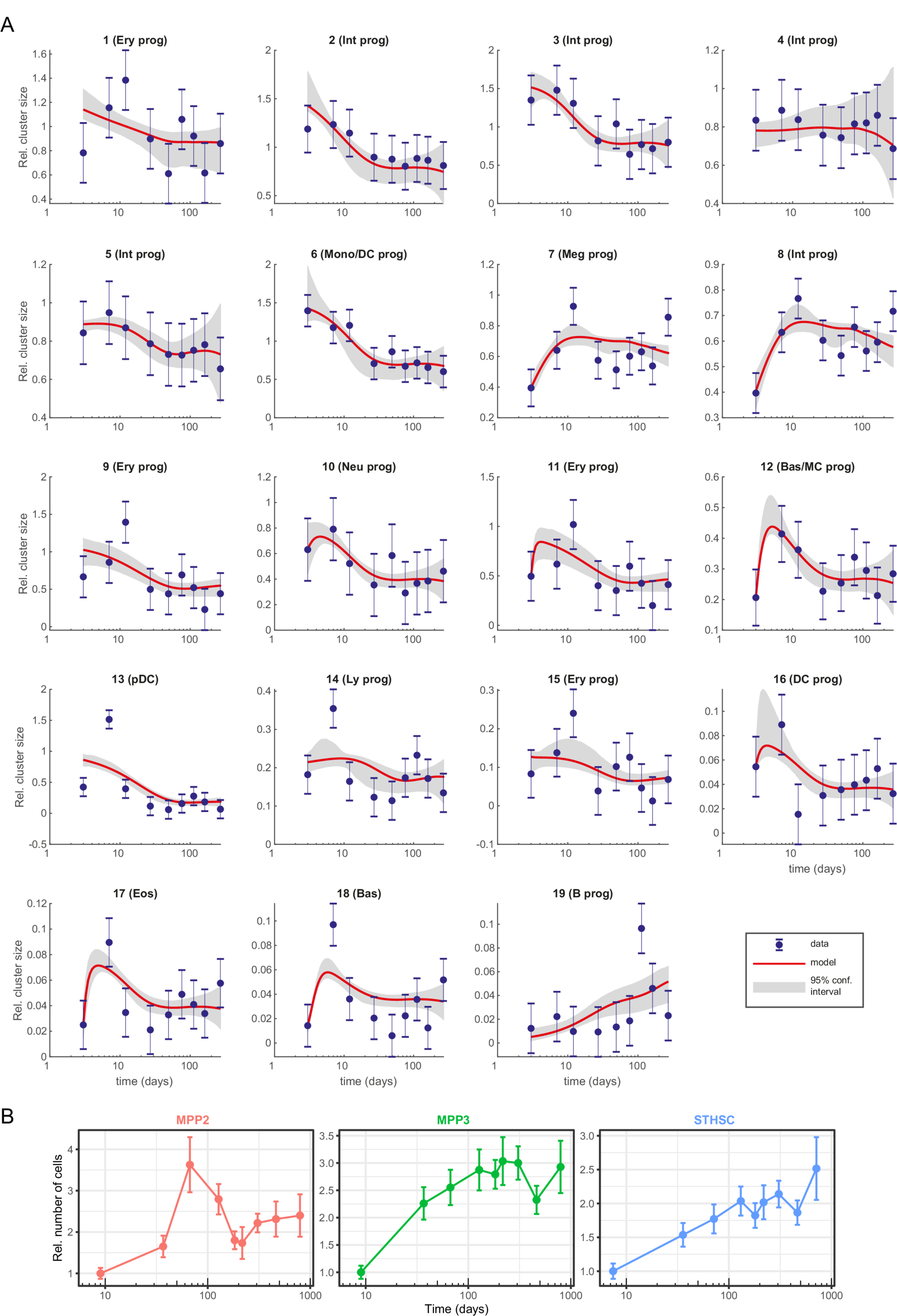
(A) Best discrete model fit (with 95% confidence intervals) for relative cluster size (normalized to cluster 0, based on Tom^-^ cells). (B) Total number of cells per mouse in the indicated populations normalized to the first time-point. Based on flow cytometry data from (Barile et al., 2020). Error bars indicate pooled standard error of the mean.

**Figure S7.**
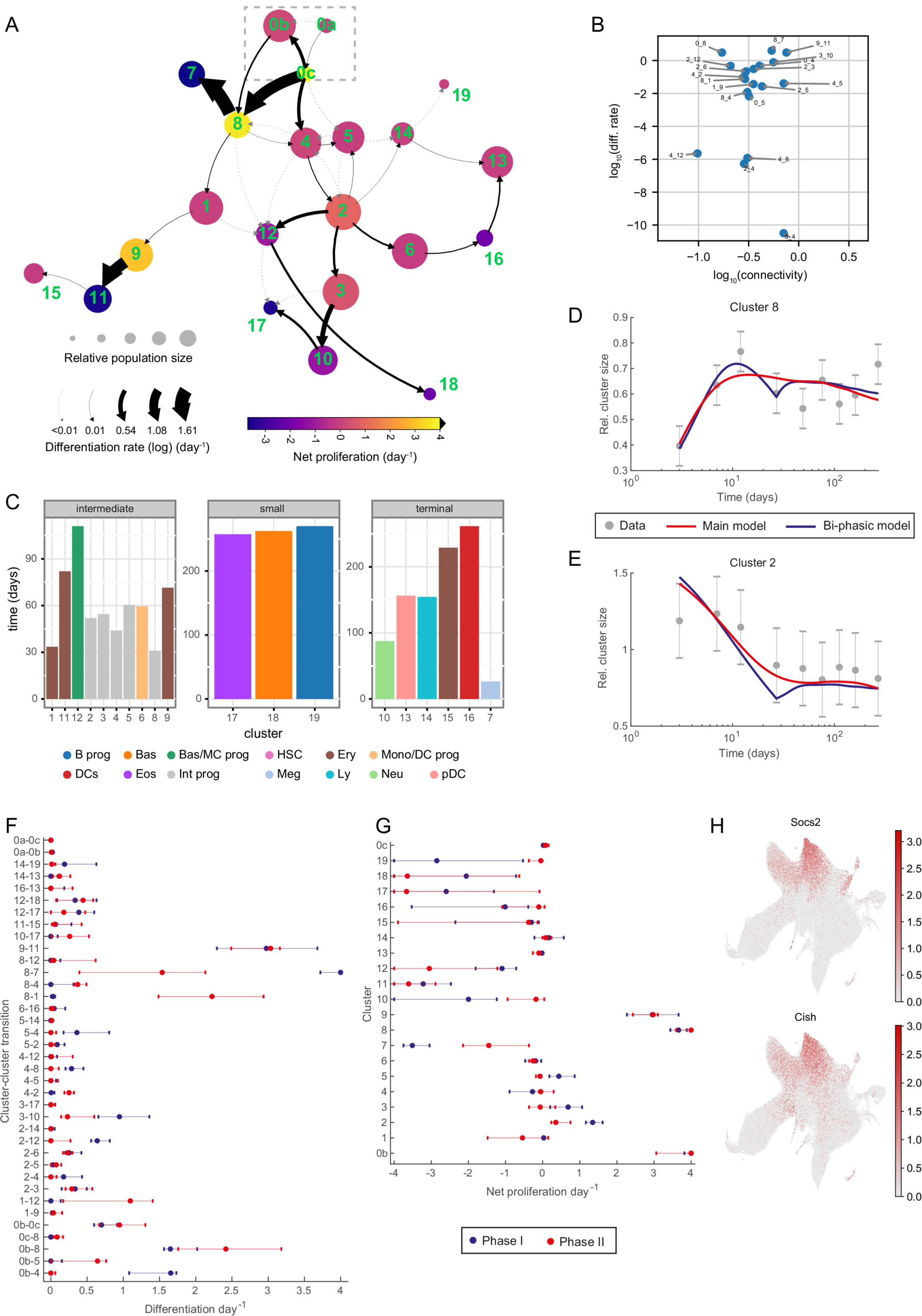
(A) Graph abstraction view of the dynamics model. Size of the nodes is proportional to square roots of relative cluster size, node color is proportional to net proliferation rate, arrows indicate differentiation directions, arrow thickness is proportional to cell differentiation rate (log-scale). (B) Scatter plot showing relation of cluster connectivity (estimated by PAGA) to differentiation rates. Only clusters 0-12 and differentiation rates greater than 10^-12^ are shown. Please note that in case of the transitions between clusters 4 and 8 two differentiation rates are plotted (each direction) (C) Related to Figure 3F, the average time required for a single cell to reach corresponding cluster when initiated in cluster 0 (journey time). (D,E) Relative cluster size (normalized to cluster 0, based on Tom^-^ cells) with the best fit for the main model (only one phase) and bi-phasic model, which permits a change in proliferation and differentiation rates after day 27. Error bars indicate pooled standard error of the mean. (F) Differentiation rates per transition for each phase of the bi-phasic model. Phase I includes the first four time-points and phase II the remaining ones. Error bars indicate 95% confidence interval. (G) Proliferation rates per cluster for each phase of the bi-phasic model. Phase I includes the first four time-points and phase II the remaining ones. Error bars indicate 95% confidence interval. (H) UMAP projection of the integrated landscape color-coded by log-normalized expression of indicated JAK/STAT target genes (Morris et al., 2018).

**Figure S8.**
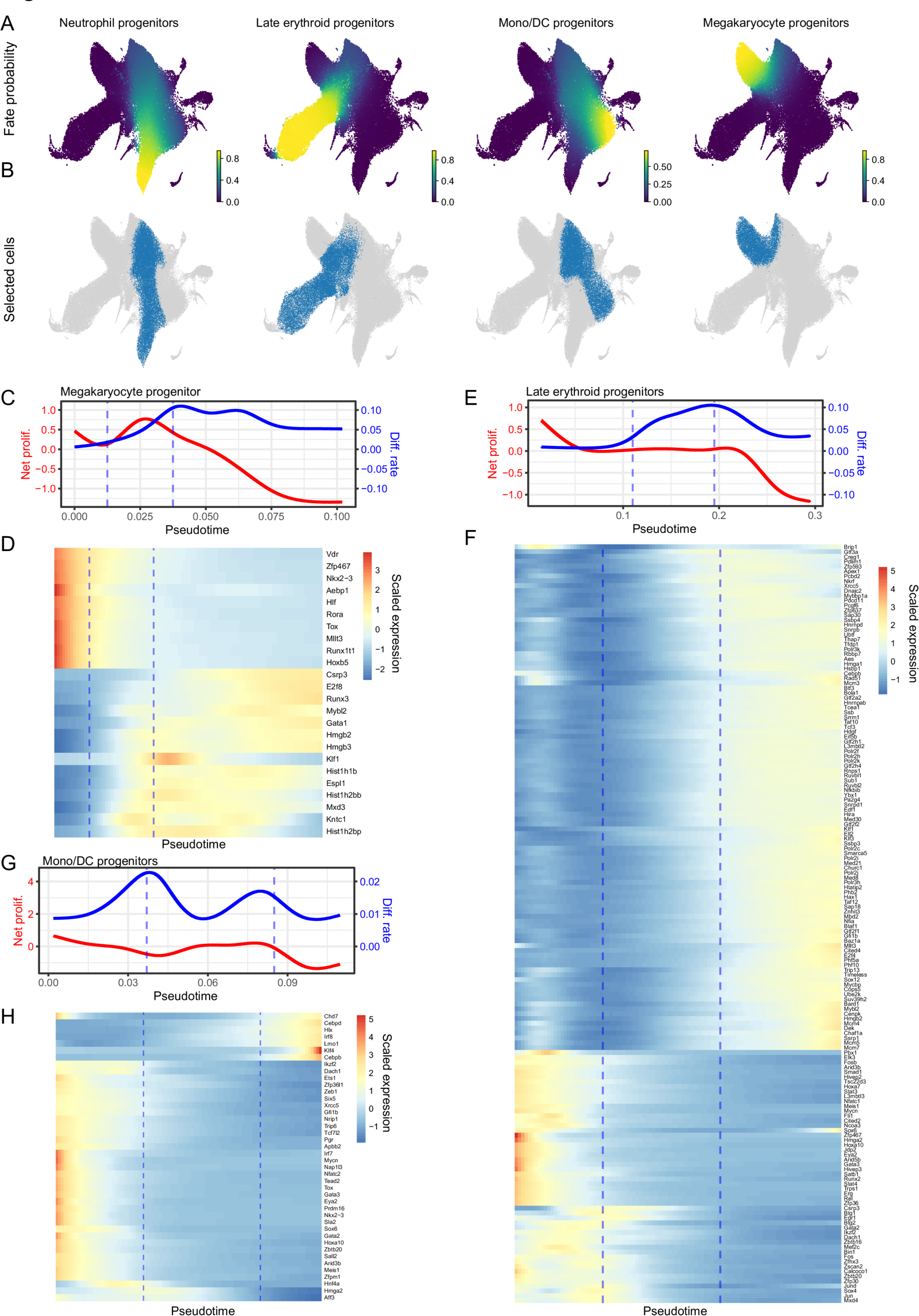
(A) UMAP projections of the HSPC landscape color-coded by cell fate probability for respective lineages (estimated with pseudotime kernel). (B) UMAP projections with cells selected for respective trajectories color-coded in blue. (C) Pseudodynamics fitted net proliferation and differentiation rate parameters along pseudotime for the megakaryocyte trajectory. Vertical lines indicate the region of interest. (D) Heatmap of TFs differentially expressed around the region of interest shown in C. (E) Pseudodynamics fitted net proliferation and differentiation rate parameters along pseudotime for the erythroid trajectory. Vertical lines indicate the region of interest. (F) Heatmap of TFs differentially expressed around the region of interest shown in E. (G) Pseudodynamics fitted net proliferation and differentiation rate parameters along pseudotime for the monocyte/dendritic cell trajectory. Vertical lines indicate the region of interest. (H) Heatmap of TFs differentially expressed around the region of interest shown in G.

**Figure S9.**
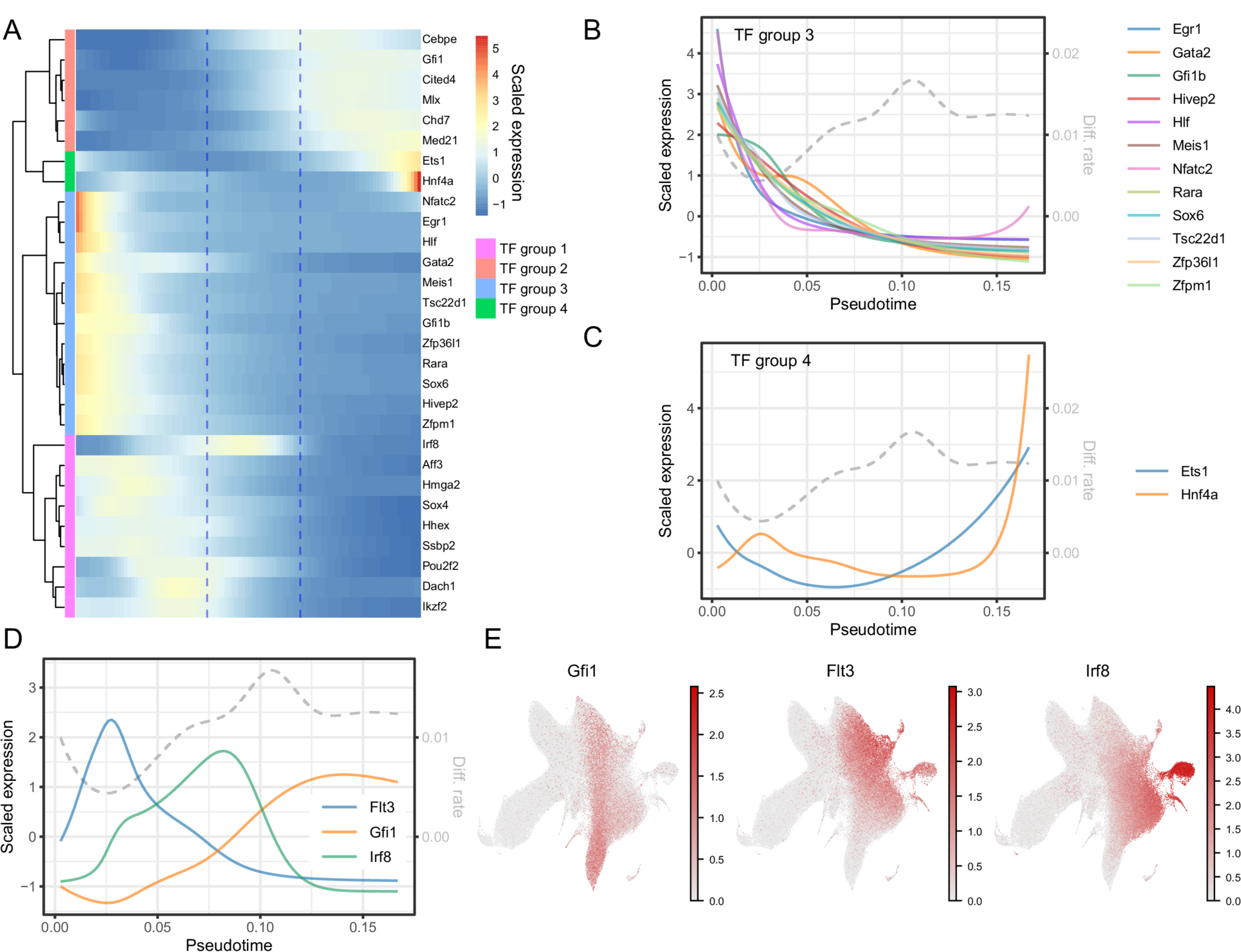
(A) Heatmap of differentially expressed TFs around the region of interest shown in Figure 4E for neutrophil trajectory. TFs are hierarchically clustered, 4 color-coded groups are plotted separately in Figures 4F and S9B, C. (B,C) Fitted gene expression values along pseudotime for neutrophil trajectory for TF groups 3 and 4, see A. (D) Fitted gene expression values along pseudotime for neutrophil trajectory for the *Gfi1*, *Flt3*, *Irf8* genes. (E) UMAP projections of the integrated HSPC landscape color-coded by log-normalized expression of genes *Gfi1*, *Flt3*, *Irf8* genes.

**Figure S10.**
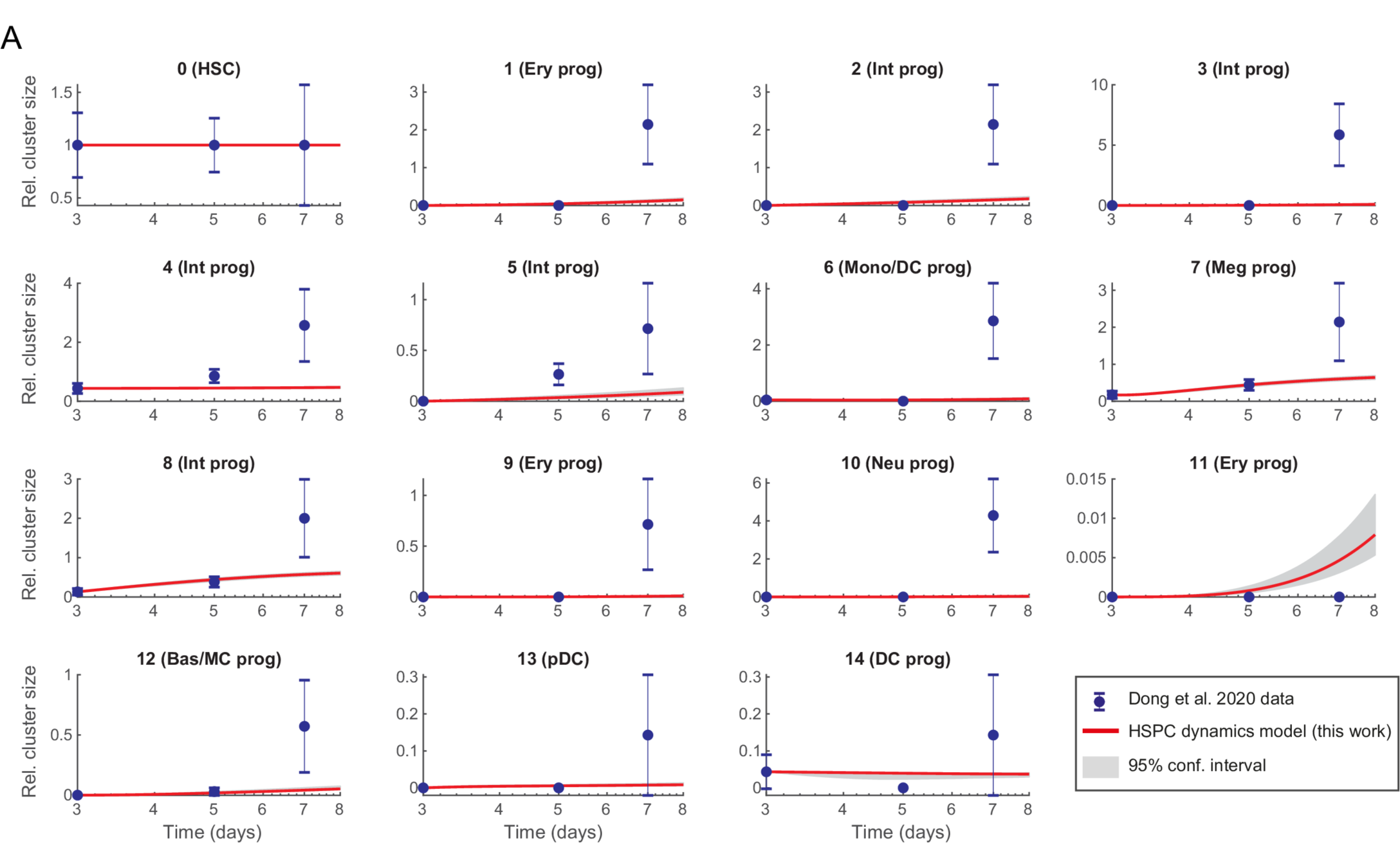
(A) Related to Figure 5. Predicted relative cluster size (red line with 95% confidence interval) based on day 3 data from (Dong et al., 2020). Observed data shown in blue. Error bars indicate propagated standard error of the mean.

**Figure S11.**
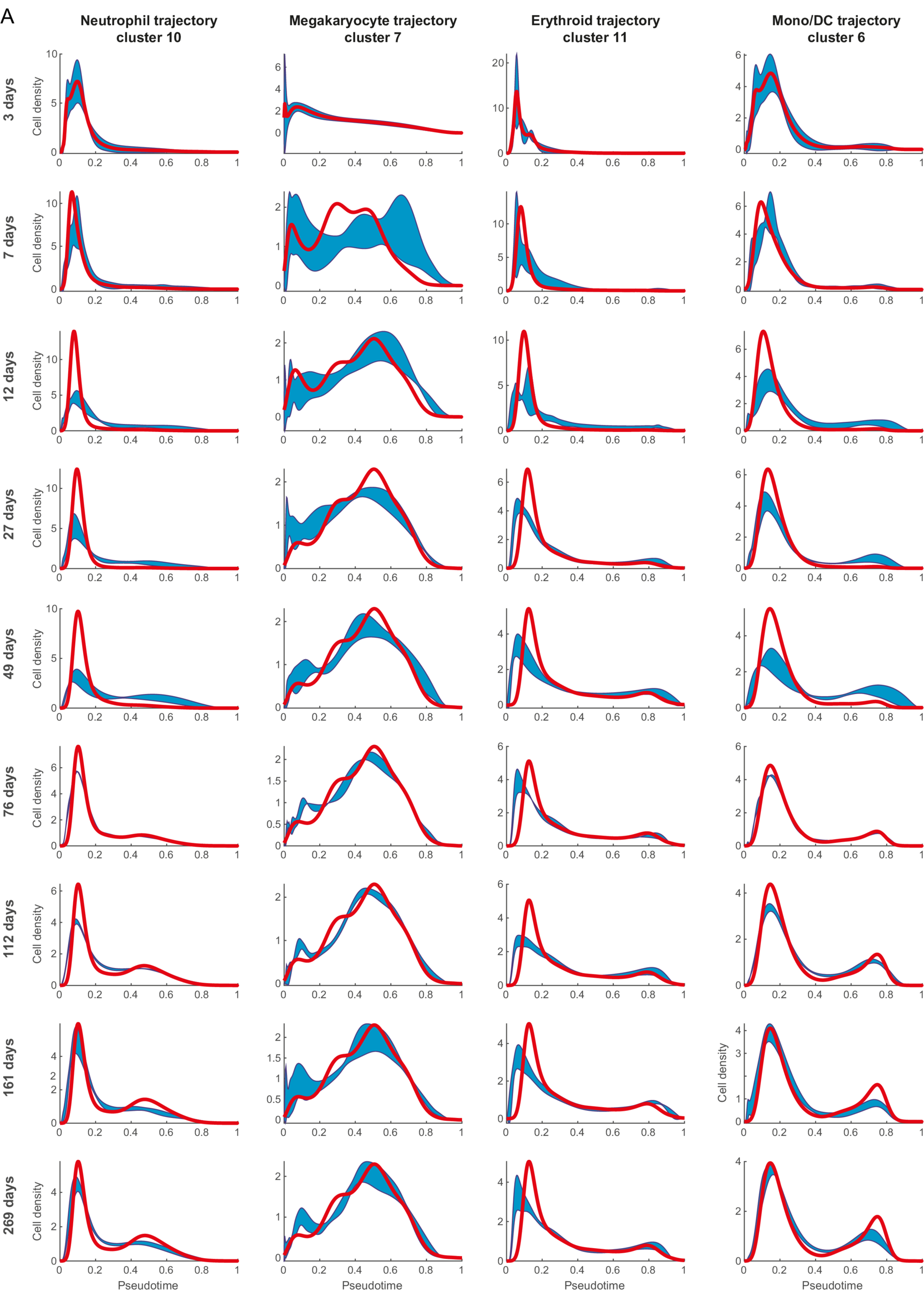
Continuous model best fits (red line) and standard deviation ranges around the mean (shaded areas) for the indicated trajectories.

## Supplementary tables legends

**Table S1.Details of the manual annotation of the integrated HSPC landscape**. Table contains the manual annotation for each cluster, indication whether the cluster was filtered out and lists of key marker genes used in the annotation process.

**Table S2. Oligonucleotide sequences.** Table containing DNA sequences of the oligonucleotides used in this work.

